# Parallel motion vision pathways in the brain of a tropical bee

**DOI:** 10.1101/2022.12.16.520712

**Authors:** Anna Honkanen, Ronja Hensgen, Kavitha Kannan, Andrea Adden, Eric Warrant, William Wcislo, Stanley Heinze

**Author notes:** These authors contributed equally (Anna Honkanen, Ronja Hensgen).

## Abstract

Spatial orientation is a prerequisite for most behaviors. In insects, the underlying neural computations take place in the central complex (CX), the brain’s navigational center. In this region different streams of sensory information converge to enable context-dependent navigational decisions. Accordingly, a variety of CX input neurons deliver information about different navigation-relevant cues. In bees, direction encoding polarized light signals converge with translational optic flow signals that are suited to encode the flight speed of the animals. The continuous integration of speed and directions in the CX can be used to generate a vector memory of the bee’s current position in space in relation to its nest, i.e. perform path integration. This process depends on specific, complex features of the optic flow encoding CX input neurons, but it is unknown how this information is derived from the visual periphery. Here, we thus aimed at gaining insight into how simple motion signals are reshaped upstream of the speed encoding CX input neurons to generate their complex features. Using electrophysiology and anatomical analyses of the halictic bees *Megalopta genalis* and *Megalopta centralis*, we identified a wide range of motion-sensitive neurons connecting the optic lobes with the central brain. While most neurons formed pathways with characteristics incompatible with CX speed neurons, we showed that one group of lobula projection neurons possess some physiological and anatomical features required to generate the visual responses of CX optic-flow encoding neurons. However, as these neurons cannot explain all features of CX speed cells, local interneurons of the central brain or alternative input cells from the optic lobe are additionally required to construct inputs with sufficient complexity to deliver speed signals suited for path integration in bees.

## Introduction

Most animals have to navigate their environment to locate food, find mates, seek shelter or to return to the safety of their nest. Using visual information to guide navigation is highly effective and thus widespread across the animal kingdom. This is because vision delivers instantaneous information about an animal’s wider surroundings - useful to recognize objects of interest, but also to estimate one’s orientation in space and the speed of one’s movements. While recognizing and assessing objects benefits from having high resolution vision (Nilsson, 2013), using vision to obtain directional information and to judge self motion is feasible with much lower resolution and thus at much lower cost for the animal (Baddeley et al., 2011, Zeil, 2012). Consistent with this idea, different pathways have evolved that process these visual tasks in parallel. Under the continuous constraints of a limited energy budget, each pathway has evolved to integrate its input over different spatial and temporal scales, optimally matching the statistics of the sensory variables to be extracted (Nassi and Callaway, 2009).

Neurally, this is manifested in segregated, parallel circuitry that extracts relevant information from photoreceptor outputs and relays it to downstream brain centers that control behavioral decisions. One set of visually guided behaviors that require such multiple parallel streams of information to be integrated are navigation and orientation strategies. While carried out by most animals, including mammals, it is insects that have proven to be a highly accessible group of animals in which to study visually guided navigation, both behaviorally and neurally (Collett, 2019, Heinze, 2017, Honkanen et al., 2019, de Jongh, 2020, Srinivasan and Zhang, 2000, Warrant and Dacke, 2016). When, for example, bees find their way back to their nest after a long and convoluted foraging flight, they can use sky compass cues and the visual panorama to determine their bearing (reviewed in Srinivasan (2015), Towne et al. (2017)), objects in the environment to identify points of interest, and optic flow patterns across their retina, generated by their own movements, to assess their speed and orientation in space (reviewed in Srinivasan and Zhang (2004), Srinivasan et al. (1999)). How these various sources of visual information are combined to produce navigation behavior remains poorly understood.

We recently identified the central complex (CX) of the bee brain as a site for convergence of visual sky compass cues and translational optic flow information (Stone et al., 2017). The CX is a group of higher order neuropils that consists of the protocerebral bridge, the fan-shaped body, the ellipsoid body and the noduli (reviewed in Pfeiffer and Homberg (2014)). As compass cues and optic flow information are suited to encode compass bearing and the speed of the animal during flight, the highly conserved CX was proposed to serve as an integrator that generates a visually based vector memory as the neural basis for path integration. While the pathway for transferring sky compass information from the eyes to the CX is well established across various insects (fruit fly: Hardcastle et al. (2021), Warren et al. (2019); monarch butterfly: Heinze et al. (2013), Heinze and Reppert (2011, 2012); locust: reviewed in el Jundi et al. (2014); dung beetles: Dacke and el Jundi (2018), el Jundi et al. (2015)), much less is known about how optic flow information reaches this region. Although responses to wide field motion have been found in the CX of several insect species, these responses are either located in intrinsic neurons of the CX (columnar polarization sensitive neurons in locusts, Rosner et al. (2019), Zittrell et al. (2022)), located in anatomically unidentified neurons (cockroach, Kathman et al. (2014)), or were generally weak, non direction-selective or state-dependent (locust, Rosner et al. (2019), Zittrell et al. (2022); flies, Weir et al. (2014)). In contrast, pronounced responses to optic flow, in particular to translational optic flow, were consistently identified in input neurons of the CX noduli (Bausenwein et al., 1994, Lu et al., 2021, Stone et al., 2017).

The neurons encoding the speed of translational optic flow in bees also specifically targeted this small CX compartment (Stone et al., 2017). The information reaching the noduli is complex and differs substantially from that encoded by optic flow sensitive lobula plate tangential neurons (LPTCs). While both encode the speed and direction of optic flow patterns, most crucially, the CX input neurons (TN cells) integrate signals from the entire panorama and are selective to horizontally expanding optic flow (Stone et al., 2017). Motion in one half of the panorama leads to inhibition, while it leads to excitation in the opposite half, a pattern that is inverted for the opposite movement direction. The four individual TN neurons that innervate the two noduli are then arranged in a way that allows the encoding of four cardinal directions of translational movements, with each principal axis being 45° offset from the bee’s body axis. Using this arrangement as a population code allows the bee to encode fully holonomic movements, i.e. movements for which the movement direction is independent of body orientation (Stone et al., 2017). This neural layout is also highly similar in the fruit fly *Drosophila* (Currier et al., 2020, Lu et al., 2021, Lyu et al., 2022, Matheson et al., 2022), suggesting that it is part of a fundamentally conserved core navigation circuit. While the input regions of the TN neurons are described in multiple species (*Megalopta*: (Stone et al., 2017), honeybee, *Apis mellifera*: (Hensgen et al., 2021), *Schistocerca*: (von Hadeln et al., 2020), *Drosophila*: (Hulse et al., 2021), it is currently not known which neurons are upstream of these cells and how the pathway between the retina and TN cells transforms simple, elemental motion signals into the complex panoramic signals that are suited to encode holonomic motion.

To delineate the visual pathways involved in processing optic flow, and to narrow down potential sources of optic flow information in the CX, we have characterized the physiology and morphology of optic lobe projection neurons, in the search for neurons that matched the physiological properties of TN neurons. As the work on the TN neurons was carried out in the nocturnal sweat bee *Megalopta genalis* (Stone et al., 2017), we have continued to explore this species. Given that more than 100 million years of evolutionary history separate these halictid bees from better described bees such as honeybees or bumblebees, our work also provides a basis for comparing visual circuits across the bee phylogeny in order to identify the fundamentally shared circuit components as well as any specializations that can be correlated to a nocturnal lifestyle in a dense jungle environment.

## Material and Methods

### Animals

Adult bees of the genus *Megalopta* (species *M. genalis* and *M. centralis*) were caught from the wild using light traps (white sheets illuminated by a bright light source containing UV wavelengths; or LepiLED (Gunnar Brehm, Jena, Germany). Traps were placed ca. 2 m above ground within small canopy openings of the tropical forest on Barro Colorado Island (field station of the Smithsonian Tropical Research Institute), located in the Panama Canal, or near Gamboa, Panama. Trappings were carried out during the activity phase of the bees between 4:00am and 5:30am in the morning, i.e. during early morning twilight. Caught bees were kept individually in 50 ml plastic vials, equipped with two cotton balls, one soaked in honey solution as well as one soaked in water. Vials were kept at room temperature in a dark secondary container (small amounts of natural light were allowed to reach the bees to ensure continuous circadian entrainment). Bees were used for experiments within two weeks after capture to ensure healthy condition. With few exceptions of males, most used bees were large to medium sized females. All experiments were conducted during the Panamanian dry season, between January and June. Collection permit: MiAmbiente Scientific permit No. SE/A-34-19, MiAmbiente export permit No. SEX/A-52-19.

### Intracellular electrophysiology

Intracellular recordings were carried out with sharp-tipped electrodes (resistance 50-150 MΩ) drawn from borosilicate glass capillaries with Sutter P-97 puller (Novato, California). Bees were anaesthetised on ice until immobile and waxed to a plastic holder. Legs and wings were removed and mandibles were immobilised with wax for increased stability of the preparation. A rectangular opening for the recording electrode was cut frontally into the head capsule. This hole covered the area flanked by the two compound eyes, antennae, and ocelli. The opening was then cleared of tracheae, fat tissue and air sacks. The brain surface was shortly exposed to Pronase (crystals applied directly), after which the neural sheath was removed with tweezers. A silver wire was placed in the rostral part of the head (near mandibles) as reference electrode. To allow optimal brain access, the preparation was placed vertically in the center of an also vertically mounted virtual reality arena (upwards in real world coordinates was hence forward from the animal’s point of view; dorsal for the animal was sideways in real world coordinates). The recording electrode (tip filled with 4% Neurobiotin (Vector Laboratories, Burlingame, California) in 1 M potassium chloride, backed up with 1 M potassium chloride) was inserted frontally under visual control via a stereo microscope, using the antennal lobes and the vertical lobes of the mushroom body as landmarks. Because of the vertically mounted preparation, we used a 90° electrode holder to lower the electrode vertically and advance its tip from anterior to posterior through the brain. The electrode was controlled with a micromanipulator in stepping mode (Sensapex, Oulu, Finland). As we were interested in visually responsive neurons that might supply the central complex (CX) with optic-flow information, we targeted areas posteriorly, ventro-laterally or dorsally of the central body. Once cells were impaled and the stimulation protocol was successfully tested, a depolarizing current (2 nA, 3 min) was applied to iontophoretically inject Neurobiotin into the recorded neuron. Recordings were performed at all times of the day on 160 female and male bees between 2014 and 2019. Of those, 37 recordings yielded the neurons presented in this paper. Signals were amplified with a BA-03X amplifier (NPI), digitized using CED-1401 micro (Cambridge Electronics Design, Cambridge, England), and recorded with Spike2 software (Cambridge Electronics Design). All recordings were performed at room temperature (ca. 25°C).

### Visual stimulation

For visual stimulation, we used a cylindrical LED arena, covering 360° of azimuthal space and 55° of vertical space (equal parts above and below the horizon). The arena was assembled from 96 commercial LED-arrays of 8×8 570 nm LEDs, mounted on FlyPanels-G3, controlled by a panels display control unit (all parts: IO-Rodeo, Pasadena, USA) and had an angular resolution of 1.5°. The LED arena was oriented at a 90° shift with respect to real world coordinates, so that upwards (real world) was forward (arena coordinates) for the animal. This panoramic LED arena was complemented by a small, UV illuminated linear polarizer (PUV 2, Spindler & Hoyer). The polarizer was driven by a Micos DT-50 rotation stage (controlled via MoCo controller; Micos) and illuminated by two UV LEDs (365 nm) that were controlled via an Arduino UNO (Arduino LLC, Somerville, Massachusetts), serving as a USB controlled switch to allow current through the LEDs from a 9 V battery. The laterally mounted polarizer covered 13.3° of the animal’s dorsal field of view. LED panels in the arena, the Arduino and the rotation stage were controlled via an integrated, custom designed software on MATLAB (MathWorks, Natick, Massachusetts).

To generate optic flow stimuli, we used sinusoidally modulated gratings of different spatial frequencies, shown at different movement speeds and directions (see below for details). To simulate optic flow experienced by the animal during yaw rotations, the gratings moved coherently around the bee in the horizontal plane in one of two directions. During clockwise grating motion (from the perspective of the bee, simulating counterclockwise yaw rotations) the grating moved towards the right in front of the animal and to the left behind the animal, and vice versa for counter-clockwise grating movements. Throughout the paper, we will refer to these simulated yaw rotations as clockwise and counter-clockwise rotational optic flow. To simulate optic flow experienced by the animal during forward or backward translational movement, we generated a simplified optic flow stimulus that captured the differential movement directions of the global optic flow pattern around the animal (identical to Stone et al. (2017)). We used the identical gratings as for simulated yaw rotations, but moved them in a front-to-back direction (progressive movement) or in a back-to-front direction (regressive movement) on both sides of the bee simultaneously, generating a point of expansion in front or directly behind the bee. We interpret progressive movement as simulating forward translation of the bee, while regressive motion simulates backward translation of the bee. While this stimulus lacks the vertical expansion and changes in spatial frequency in different parts of the visual field that are both features of translational optic flow, it matches the flow fields generated by translational movements in wide areas of the lateral fields of view, thus serving as an approximation of translational optic flow.

Optic-flow stimuli were shown either in fixed order or in random sequences of individual stimulus bouts separated by darkness. In all cases, each bout consisted of 0.5 s stationary display of the stimulus pattern followed by 3 s of movement at constant velocity, followed by 3 s of darkness. Progressive and regressive motion stimuli were presented with fixed spatial frequencies of 0.067 cycles/° and movement velocities ranging from 15°/s to 120°/s. Clockwise and counter-clockwise yaw rotations were presented either with fixed spatial frequencies of 0.067 cycles/°combined with velocities ranging from 10°/s to 200°/s, or with fixed velocities of 60°/s combined with varying spatial frequencies of 0.033 cycles/°, 0.067 cycles/°, 0.1 cycles/°, and 0.133 cycles/°. Accordingly, temporal frequencies (as the product of spatial frequency and velocity) of progressive and regressive motion stimuli ranged from 1.01 cycles/s to 8.04 cycles/s, while clockwise and counter-clockwise yaw rotations had temporal frequencies of 0.67 cycles/s to 13.4 cycles/s.

Receptive fields were mapped using a narrow vertical stripe (width: 7.5° full arena height) that moved around the entire panorama at constant speed (60°/s) either clockwise or counter-clockwise. Each stimulus consisted of two clockwise rotations followed by two counter-clockwise rotations. The bar was introduced into the arena behind the bee and remained stationary for 0.5 s before movement commenced. Similarly, in some recordings we applied a 7.5° wide horizontal bar (wrapping around the arena) to map local responses to vertical motion. Control voltages were recorded for all stimuli, indicating the timing of displayed frames in the virtual reality arena and the angular position of the rotation stage controlling the polarizer.

As *Megalopta* bees are nocturnal, we also tested responses to dim-light stimuli by introducing one to three layers of ND0.9 neutral density filters into the LED arena (LEE 211 0.9ND, LEE Filters Worldwide, Andover, UK). As seeing any clear responses to any of the stimuli with three ND filters was very rare, but weak responses were frequently seen with two filters, we focused all analysis and stimulation to conditions with two filters or fewer. Fractional transmittance of 1, 2, and 3 layers of ND0.9 filters is ca. 12.5%, 1.56%, and 0.19%, respectively. The illuminances were measured with Hagner digital luxmeter EC1 (Hagner, Solna, Sweden) at the center of the arena for a moving sinusoidal stimulus as 24.8, 3.4, and 0.4 lx for 0, 1 and 2 layers of ND filters, respectively. The readings for the two darker conditions correspond with the expected values of 3.1 and 0.39 lx. The filter film was fixed into three nested cylinders that fit tightly inside the arena allowing the removal of either just the innermost layer or more layers at the time. While experiments in 2014 and 2015 were performed without these filters, all remaining experiments were carried out at reduced light levels. In these experiments we started the stimulus presentation with the dimmest light intensity we were going to use (e.g. two layers of ND filter) and let the bee dark adapt to that intensity. After testing all stimuli, we progressed to take filters out one by one and repeating the stimuli over the course of the recording. Experiments were usually started with two ND filters and proceeded to one and finally no filters. As no qualitative difference in the responses between different light levels were observed across our recordings, and not all light levels were tested in all neurons, we did not include light level as a parameter for analysis in this paper. For displayed data, we used either no filters or 2 ND filters. If not otherwise stated, the default of 2 ND filters was used to obtain the recordings.

Polarized light stimuli were presented by switching on the UV LED and subsequently rotating the polarizer by 360° in clockwise and counter-clockwise directions. A control voltage was recorded that corresponded to the position of the rotation stage. No neuron presented in this study responded to this stimulus.

When describing receptive fields and stimulus positions, the terms right and left are always used from the perspective of the bee.

### Neurobiotin Histology and Immunohistochemistry

Neurobiotin injected brains were processed as follows. Injected brains were removed from the head capsule and fixed in neurobiotin fixative (4% paraformaldehyde, 2% saturated picric acid, 0.25% glutaraldehyde) overnight at 4°C. Brains were transferred to 0.1 M PBS until further processing. After rinsing the brains for 4 × 15 min in PBS, they were incubated with Streptavidin conjugated to Cy3 (1:1000, in 0.1 M PBT (PBS plus 0.3% TritonX-100)) for 3 days. The brains were then washed (4 × 20 min PBT, 2 × 20 min PBS) and dehydrated in an increasing ethanol series. Finally, they were cleared in Methyl salicylate and mounted in Permount (between two coverslips, separated by spacers). The brain samples were taken to Lund University, Sweden, for microscopy and 3D reconstructions of the cells.

Staining for synapsin and serotonin on thick sections followed the protocol described by Adden et al. (2020). Brains were dissected out of the head capsule in moth ringer solution (150 mM NaCl, 3 mM KCl, 10 mM TES, 25 mM sucrose, 3 mM CaCl_2_; based on King et al. (2000)) and fixated in Zambonis fixative (4% PFA, 7.5% picrid acid in 0.1 M phosphate buffer) overnight at 4°C. The brains were then washed in PBS (3 × 10 min), embedded in albumin-gelatin (4.8% gelatin and 12% ovalbumin in demineralized water) and postfixed in 4% formaldehyde solution in 0.1 M PBS overnight at 4°C. The brains were cut into 40-,tm thick sections using a vibrating blade microtome (Leica VT1000 S, Leica Biosystems, Nussloch, Germany). The sections were washed in 0.1 M PBS (3 × 10 min) and pre-incubated in 5% NGS in 0.1 M PBT for 1 hr. The primary antibody solution (1:50 anti-synapsin, 1:1000 anti-5HT, 1% NGS in PBT) was applied overnight at room temperature. After rinses in PBT (8 × 10 min) the sections were incubated in the secondary antibody solution (1:300 GAM-Cy5, 1:300 GAR-488, 1% NGS in PBT) overnight at room temperature. Following rinses in PBT (3 × 10 min), the sections were mounted on chromalaun/gelatin-coated glass slides and left to dry for at least 5 hr. The mounted sections were dehydrated in an increasing ethanol series (demineralized water, 5 min; 50%, 70%, 90%, 95%, 2 × 100%, 3 min each), cleared in xylene (2 × 5 min), and embedded in Entellan (EMS, Hatfield, PA).

### Confocal microscopy, image processing and 3D reconstructions

Confocal imaging of single neuron morphologies was carried out either with a Zeiss LSM 510 equipped with a 25x long distance objective (LD LCI Plan-Apochromat 25x/0.8 Imm Corr DIC, Zeiss) or with a Leica SP8 equipped with a 20x long distance objective (HC PL APO 20Œ/0.75 Imm Corr CS2). In the Leica microscope, the 552 nm laser line was used for excitation, while in the Zeiss LSM we used the 561 nm line. For the Leica microscope, the hybrid detector (HyD) was used to maximize photon catch, either in counting mode or in standard mode with smart gain set to the lowest possible value (10%). With line accumulation set to 2 or 3, laser intensity was set to the minimum value that yielded sufficiently bright images. Neurons were imaged at a voxel size of 0.3 × 0.3 × 0.88 *µ*m or slightly larger. If several image stacks were required to cover the full extent of any one cell, these were aligned to a common reference frame (using Amira or FIJI) and used as input to the skeletonize plugin of Amira (Evers et al., 2005, Schmitt et al., 2004). Neurons were traced manually and the resulting skeletons were finalized by automatic midline fitting and diameter adjustment (using local brightness information of the image data).

When describing brain anatomy, the terms right and left are always used from the perspective of the bee.

### Data analysis

Action potentials were extracted from the recorded voltage traces using threshold based event detection in MATLAB. Only recordings with stable baseline were evaluated. Timing of events was then correlated to the recorded stimulus traces by custom designed analysis scripts. For all presented neuron types, numbers of recorded cells can be found in the results section. Within each cell type the number of cells always equals the number of bees used.

For optic-flow stimuli each stimulus bout was analyzed independently. Spikes were counted in bins of 0.25 s during stimulation intervals and the resulting mean frequencies were plotted for display of individual stimulus responses. To calculate tuning curves, the last 2 s of each stimulus interval was used to compute the mean response frequency for the analyzed condition and plotted against either stimulus velocity or spatial frequency. Background activity of a neuron was calculated as the mean activity during 2 s before the onset of the first stimulus.

Receptive field mapping was analyzed by binning each full movement of the bar around the bee in 8° bins, and counting the number of events during each bin. The result was divided by the duration of the bin to transform spike count into frequency. As clockwise and counter-clockwise movement direction was performed twice per stimulus bout, the mean and standard deviation was calculated for each bin. To transform the frequency bins from the time domain to azimuth angles, each bin was assigned an azimuth value (0-360° for clockwise direction and 360-0° for counter-clockwise direction). To plot both directions on the same linear axis, the counter-clockwise values were displayed in inverted order. For display of individual neurons’ receptive fields, the result of the averaging was low pass filtered in Matlab (3rd-order Savitzky-Golay (polynomial) smoothing filter; window-size: 7 bins). For comparing multiple recordings within a cell type, filtered data were normalized to peak frequency and averaged across neurons.

The mapping of receptive fields in the elevation domain was performed analogous to the mapping along the horizon.

## Results

### Optic lobe layout and internal structure

To enable us to efficiently compare single neurons of the *Megalopta* optic lobe across individual samples we first described its general layout and internal structure using immunohistochemistry (Fig. 1). From distal to medial, the optic lobe contained three main neuropils: the lamina, the medulla, and the lobula complex. Both the medulla and the lobula could be divided into an inner and outer subregion, based on externally visible crevices on the neuropil surfaces. The outer and inner medullae were separated by the serpentine layer, which is formed by the axons of large medulla tangential neurons (Fischbach and Dittrich, 1989). Using combined synapsin and serotonin labeling we distinguished different layers within each neuropil. These layers (also referred to as strata; Ito et al. (2014)) were visible within each neuropil. They result from the distinct organization of terminals of the neuron types that constitute the optic lobes and are apparent as differences in staining intensity and structure. We used these layers as a scaffold to localize projections of individually labeled neurons.

**Figure 1:**
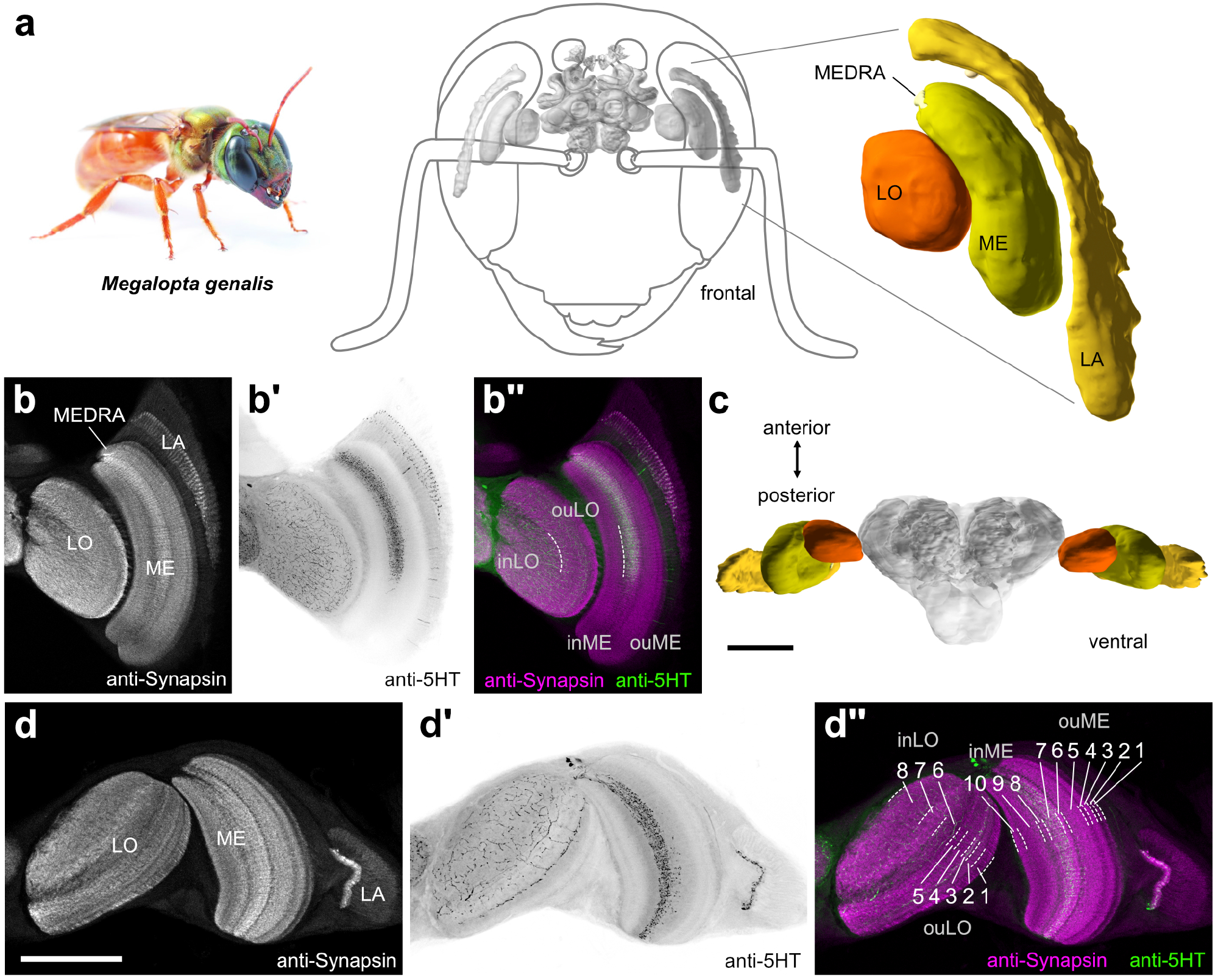
Anatomical organization of the optic lobes of *Megalopta genalis*. (a) Left: photograph of female *Megalopta genalis* (with permission from Ajay Narendra). Anterior view of the 3D reconstruction of the bee brain (gray) embedded in a schematic illustration of the bee’s head. Optic lobe (colour) of the left hemisphere enlarged on the right; lobula (LO), medulla (ME), dorsal rim of the ME (MEDRA), lamina (LA). (b-b”) Single optical sections of frontal vibratome sections stained against synapsin (gray in (b), magenta in (b”)) and serotonin (5HT, black in (b’), green in (b”)).(b”) Both LO and ME can be divided into an inner region (inLO, inME, respectively) and an outer region (ouLO, ouME, respectively). (c) Ventral view of the 3D reconstruction of the *Megalopta* brain with highlighted optic lobes. (d-d”) Single optical sections of horizontal vibratome sections stained against synapsin (gray in (d), magenta in (d”)) and 5HT (black in (d’) and green in (d”)). (d) The inLO and ouLO consist of three and five layers, respectively. The inME and ouME consist of three layers and seven layers, respectively. Scale bars = 500 *µ*m (c), 200 *µ*m (d)

The lamina was the neuropil with the least pronounced internal structure. While it appeared mostly homogeneous, a narrow strip of tissue at the proximal end of the lamina stood out with very bright synapsin immunoreactivity (Fig. 1b,d). This layer was also the only part of the lamina with serotonin immunoreactive fibers (Fig. 1b’,d’). In the second optic neuropil, the medulla, we distinguished ten layers (Fig. 1d”) and an additional, distinct dorsal region without obvious internal structure, the dorsal rim medulla. This region receives photoreceptor terminals originating in the polarization sensitive region of the compound eye, the dorsal rim area (DRA). Of the ten identified medulla layers 1-7 were located in the outer medulla, and layers 8-10 were located in the inner medulla (Fig. 1b”,d”). Layer 7, close to the boundary of inner and outer medulla, was characterized by a dense mesh of serotonergic fibers, which extended from the dorsal edge of the medulla to the level at the eyes equator (Fig. 1b’). As in other hymenopteran insects, the third optic neuropil, the lobula complex, consisted of only one neuropil, the lobula. This region comprised eight distinct layers, of which layers 1-5 formed the outer lobula and layers 6-8 the inner lobula (Fig. 1b”,d”). Serotonin immunoreactivity was particularly strong in layer 3, while a loose mesh of serotonin-positive processes permeated the largest parts of the inner lobula (layers 6-8) with some fibers penetrating into the proximal layers of the outer lobula (Fig. 1b’,b”,d’,d”).

### Single cell recordings

In total, we have recorded from 37 neurons with projections in the *Megalopta* optic lobe. These belonged to 18 morphologically distinct cell types that we divided into four categories based on their branching patterns. These categories comprise lobula projection neurons (LO-PNs; 7 types; 13 recordings), inter-medulla neurons with central brain projections (ME-ME-PNs; 5 types; 14 recordings), inter-medulla/lobula neurons (MELO-MELO-PN; 1 type; 3 recordings), and centrifugal feedback neurons to either lobula (LO-CNs; 2 types, 2 recordings) or medulla (ME-CNs; 3 types, 3 recordings). While many cell types were only found once, several types of lobula and medulla projection cells were encountered in repeated recordings.

Physiologically, all neurons responded to visual cues provided via a 360° azimuth LED arena, equipped with green LEDs (570 nm). With the exception of most centrifugal neurons, optic flow stimuli elicited strong responses and receptive fields could be mapped with a narrow, vertical bar. The cells were functionally classified according to direction selectivity and the extent and location of receptive fields. Polarized light stimuli (rotating linear polarizer, illuminated by UV-LEDs) were tested in most recordings, but did not elicit a response in any of the tested cells. This is consistent with the finding that none of the identified neurons had arborizations in the dorsal rim medulla.

In the following we will describe the different morphological neuron classes and highlight physiological response characteristics within each group of cells, aiming at delineating general patterns of information transfer between the different optic lobe neuropils and regions of the central brain.

### Lobula projection neurons

#### Morphology

We found seven anatomically distinct types of large-field lobula cells projecting from wide areas of specific layers of the lobula to the posterior protocerebrum (Fig. 2). Five types projected to bilateral regions in the protocerebrum, while the remaining two types projected either ipsilaterally or exclusively contralaterally. For all cells in which the soma was labeled, it was located near the optic stalk, either on the ventral or the dorsal side. The terminal fibers located in the lobula were fine and non-blebbed, which, together with the location of the soma, indicated that these cells receive input in this region. These arborization trees were always restricted to thin layers of the outer regions of the lobula. Within the innervated layer, most neurons extended either through dorsal regions (two types), ventral regions (one type), or medial portions of the lobula (one type), while three types innervated an entire layer. Overall, projections in the lobula were restricted to the outer lobula, with most cells innervating layer 5 (LO-PN-bilat-1,-2,-3 and LO-PN-contra-1). Two cell types possessed fibers in layers 3/4 (LO-PN-bilat-4 and LO-PN-ipsi-1), while LO-PN-bilat-5 innervated layer 2. The innervation of distinct lobula layers suggests that those cell types receive inputs from different sets of upstream neurons, a notion that was at least in part supported by distinct physiological differences.

**Figure 2:**
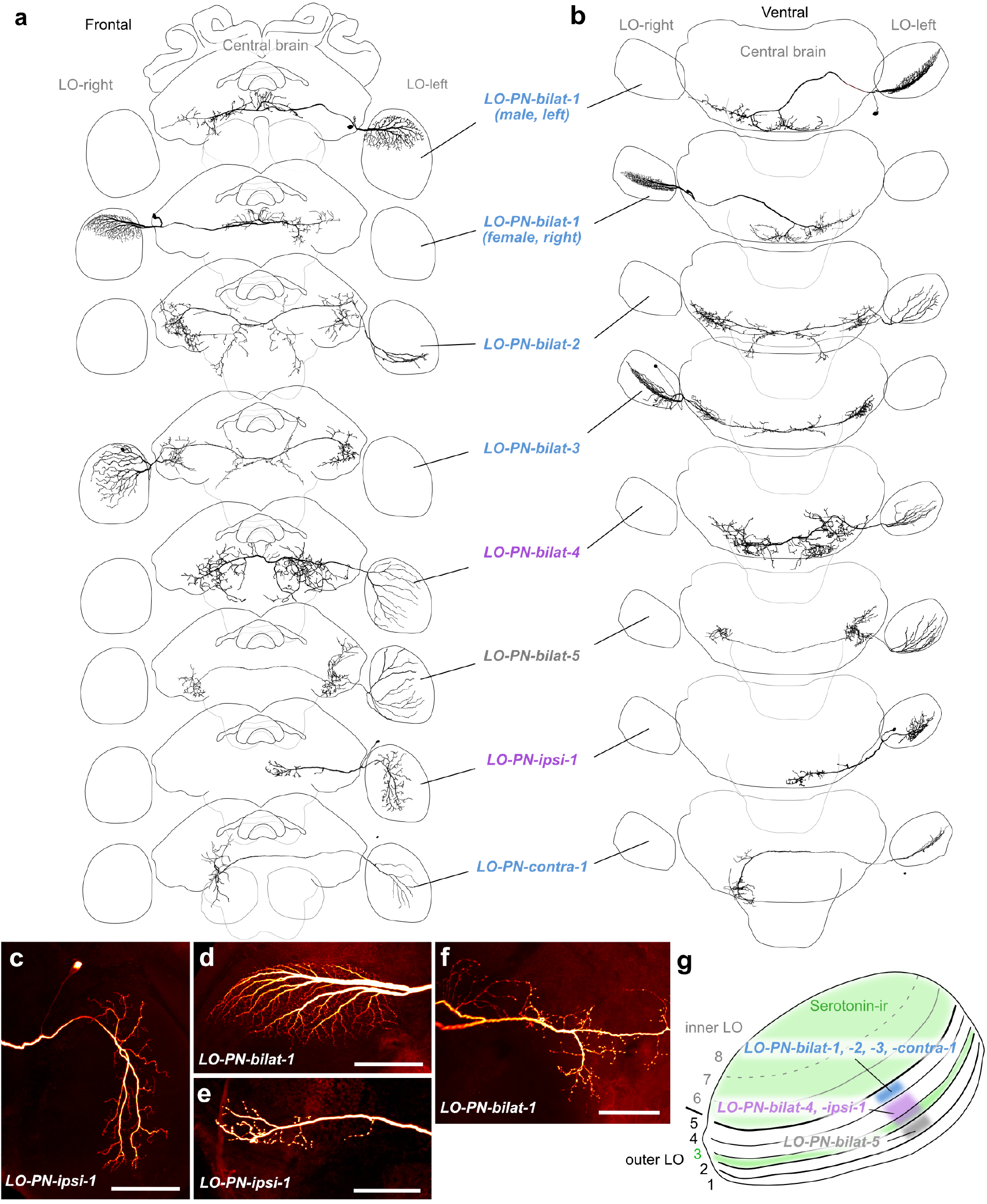
Morphology of large-field lobula projection neurons (LO-PNs). (a) 3D reconstructions of different LO-PN types (frontal view), with schematic outline of the brain for orientation. (b) Ventral view. (c-f) Confocal images of a neurobiotin-injected LO-PN-ipsi-1 neuron (c,e) and a neurobiotin-injected LO-PN-bilat-1 neuron (d,f). Maximal intensity projections of arborizations in the lobula (c,d) and the posterior protocerebrum (e,f). (g) Schematic representation of the lobula (horizontal view) illustrating the layers innervated by the different types of LO-PNs. Serotonin-immunoreactivity (ir) is distributed throughout the layers of the inner lobula (layers 6-8) and layer 3 of the outer lobula. LO-PN-bilat-1,-2,-3, and contra-1 neurons: layer 5; LO-PN-bilat-4, and ipsi-1 neurons: layers 3 and 4; LO-PN-bilat-5 neurons: layer 2. Note that the colored rectangles indicate the identity of the arborization layer, not the extent of arborizations within layers. Scale bars = 100 *µ*m

Within the central brain, all arborizations were concentrated in posterior brain regions, starting at the level of the central body, but mostly lining the posterior surface of the brain, including regions of the ventrolateral and ventromedial protocerebrum, as well as the peri-esophageal and sub-esophageal neuropils. These fibers had a strongly beaded appearance and hence were classified as output sites (Fig. 2e,f). The first cell type was encountered most frequently (n=7) and belonged to the bilateral projection neurons (LO-PN-bilat-1). It showed a consistent morphology irrespective of whether the neuron was recorded from males or females. Neurons from the right and left side of the brain possessed mirror symmetrical characteristics. All cells innervated a single layer of the outer lobula (layer 5), extending across circa one third of the neuropil, between the dorsal edge and the equator (Fig. 2d). The main neurite leaves the optic lobe posteriorly, turns anteriorly in the central brain towards the level of the mushroom body pedunculus and after passing by just ventrally of the noduli of the CX it projects towards the posterior brain surface. After crossing the midline, it branches into two major fibers, of which one continues the path towards the contralateral posterior lateral protocerebrum, where it gives rise to beaded arborizations (Fig. 2f). The second branch projects back across the midline, forming a fan of beaded terminals lining the posterior surface of the brain on either side of the midline (Fig. 2f).

Two more types of bilateral neurons (LO-PN-bilat-2, n=1, and LO-PN-bilat-3, n=1) shared a highly similar morphology with one another (Fig. 2a,b). However, LO-PN-bilat-2 neurons innervated the ventral half of the lobula, whereas LO-PN-bilat-3 neurons innervated the dorsal two thirds. Both neurons formed dense, beaded arborizations in the ventrolateral protocerebrum, approximately at the depth of the central body. The main neurite passes the midline between the esophagus foramen and the protocerebral bridge (PB). In the cell receiving input from the dorsal lobula, a sparse set of fibers were found in posterior neuropils around the esophagus. These fibers were more numerous in the cell that received input in the ventral medulla.

The fourth and fifth type of bilateral neuron (LO-PN-bilat-4 and LO-PN-bilat-5) both innervated a complete layer of the lobula but differed in their arborizations in the central brain (Fig. 2a,b). The first type projected to large regions on either side of the esophagus, extending substantially into the sub-esophageal zone. The second type formed beaded arborizations in the ventrolateral protocerebrum of both hemispheres. Similar to the other cell types, these cells also showed a pronounced polarity with beaded fibers in the central brain and smooth fibers in the lobula.

Two types of unilateral lobula projection neurons were identified, one with ipsilateral and one with contralateral central brain projections (LO-PN-ipsi-1 and LO-PN-contra-1, respectively; Fig. 2a,b). The ipsilateral cell innervated the posterior half of the lobula (Fig. 2c) and formed beaded arborizations in ventromedial neuropils (Fig. 2e). The contralateral cell extended across one half of the lobula around the equator. The main neurite of the cell passed the midline and gave rise to arborizations in ventromedial neuropils.

#### Physiology

With the exception of the LO-PN-bilat-5 neuron, we were able to physiologically characterize all morphologically described lobula neurons. Two principle types of stimuli were tested: First, a bright vertical bar moving around the bee at 60°/s (tested in all recordings; Fig.3), and second, optic flow generated by sinusoidally modulated gratings moving at constant speed, either clockwise or counter-clockwise to simulate yaw rotations, or progressive and regressive to simulate forward and backward movement, respectively (all recordings except LO-PN-bilat-2; Fig. 4). Across all recorded LO-PNs the responses to the vertical bar stimulus enabled us to distinguish two overall groups of neurons. The first one (LO-PN-bilat-1, -ipsi-1, -contra-1) was characterized by very strong responses, reaching sustained peak activities exceeding 100 impulses per second in most recordings (Fig. 3c,d bottom). The second group (LO-PN-bilat-2,-3,-4) showed more diverse and generally weaker responses (Fig. 3d, top three panels). Both groups shared pronounced direction selectivity, with strong excitation in response to the preferred direction of movement and either inhibition or no response in the anti-preferred movement direction. The only exception to this was the LO-PN-bilat-2 neuron, which showed similar responses to both movement directions (Fig. 3d, top).

**Figure 3:**
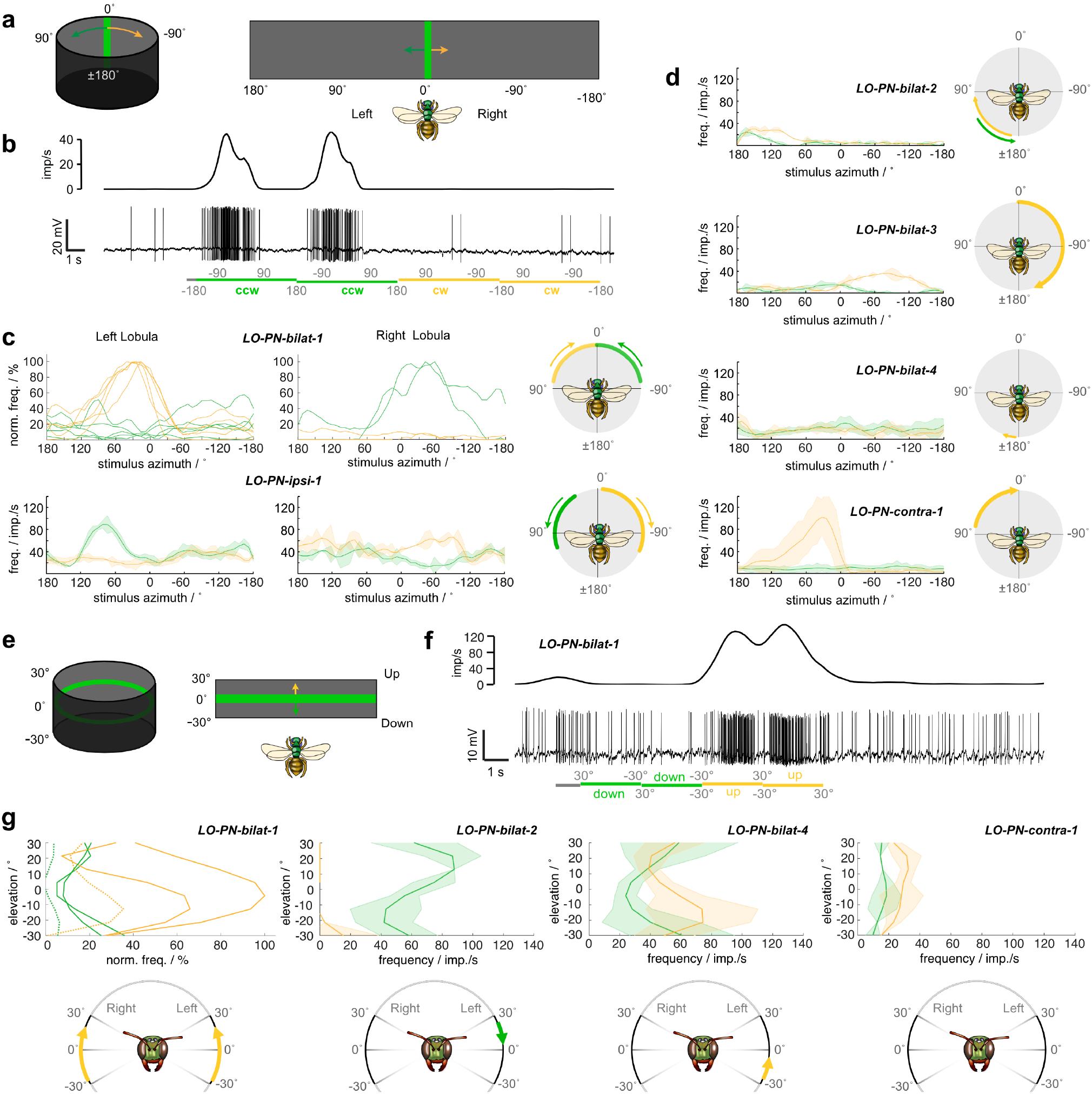
Receptive field properties of large-field lobula projection neurons (LO-PNs). (a) Left: Schematic illustration of the stimulus for mapping the horizontal receptive field. A green vertical stripe was moved around the bee in either clockwise or counter-clockwise direction at constant speed. Right: Schematic display of the flattened arena used for the graphs in (c) and (d). (b) Example response of a LO-PN-bilat-1 cell to the moving stripe. Bottom: Spike train. Top: Sliding average. Green bars, counter-clockwise (ccw); yellow bars, clockwise (cw) direction. (c) Top: Normalized average response curves of LO-PN-bilat-1 neurons of the left (n=5, no ND filter for n=3) and right (n=2, no ND filter for n=1) lobula during receptive-field mapping. Bottom: Average response curves of LO-PN-ipsi-1 neurons of the left and right lobula. Right: no ND filter. Yellow traces: cw movement, green traces: ccw movement. Cartoons on the right illustrate the approximate horizontal extent of the receptive field and the preferred movement direction of each neuron type. (d) Same as in (c, bottom) but for LO-PN-bilat-2, LO-PN-bilat-3, LO-PN-bilat-4, and LO-PN-contra-1 neurons of only one hemisphere. (e) Left: Schematic illustration of the stimulus for mapping the vertical receptive field. A green horizontal stripe spanning the whole panorama was moved up or down at constant speed. Right: Schematic display of the flattened arena used for the graphs in (g). (legend continued on next page) (f) Example response of a LO-PN-bilat-1 cell to a vertically moving stripe. Bottom: Spike train. Top: Sliding average. Green, downwards motion; yellow, upwards motion. (g) Left: Averaged responses of two LO-PN-bilat-1 neurons from the left lobula (solid lines) and one LO-PN-bilat-1 neuron from the right lobula (dotted lines) to upwards (yellow) or downwards (green) bar motion. Averaged responses of a LO-PN-bilat-2, a LO-PN-bilat-4, and a LO-PN-contra-1 cell to the same stimulus. Cartoons at the bottom, approximate vertical extent of the receptive field and the preferred movement direction of each neuron type.

**Figure 4:**
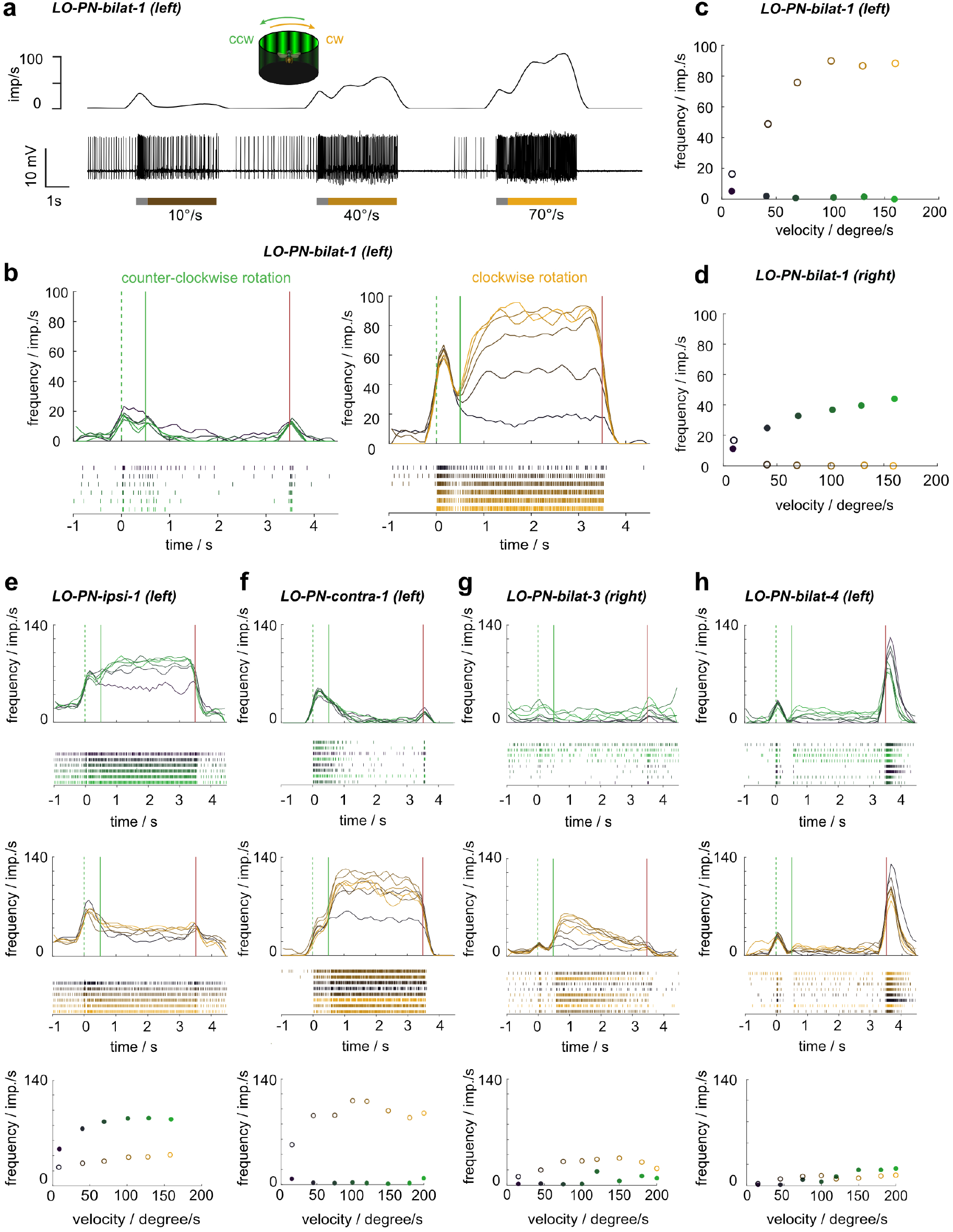
Tuning of large-field lobula projection neurons (LO-PNs) to simulated yaw rotations. (a) Example response of a LO-PN-bilat-1 cell to different velocities of clockwise rotational optic flow. Bottom: Spike train. Top: Sliding average. Gray portion of the bars: stationary display of the grating; colored portion of the bars: movement of the grating at constant speed. (b) Activity of LO-PN-bilat-1 neurons of the left lobula in response to counter-clockwise (left) and clockwise (right) rotational optic flow of different velocities (10°/s to 160°/s [dark to light green/yellow]; values: 10°/s, 40°/s, 70°/s, 100°/s, 130°/s, and 160°/s). Green dotted lines: grating presented; green solid lines: motion onset; red lines: motion stop. (c) Mean response frequency during the final 2 s of each stimulus bout shown in (b). (d) Same as in (c) but for responses of LO-PN-bilat-1 neurons of the right lobula. (e-h) Same as in (b,c) but for a LO-PN-ipsi-1, a LO-PN-contra-1, a LO-PN-bilat-3, and a LO-PN-bilat-4 cell. No ND filter in (a-e)

The small azimuthal extent of the vertical bar stimulus enabled us to test responses to localized motion stimuli, i.e. to map the receptive fields of the recorded neurons (Fig. 3a-d). The regions of azimuthal space the neurons responded to were always centered on the side of the brain containing the lobula arborizations, confirming the role of these fibers are input sites. In all neurons of group 1 (LO-PN-bilat-1, LO-PN-ipsi-1, LO-PN-contra-1) the half-width of the receptive fields along the horizon was approximately 60-90°. These regions were adjacent to the midline and in some cases extended slightly into the contralateral field of view (Fig. 3c). In contrast, group 2 neurons were more diverse; LO-PN-bilat-3 cells had a receptive field covering almost an entire hemisphere (Fig. 3d, second panel), while the not direction-selective LO-PN-bilat-2 neuron possessed a receptive field in the posterior field of view (Fig. 3d, top), consistent with its lobula fibers located on the opposite side of the lobula compared to the other neurons. The LO-PN-bilat-4 neuron showed no pronounced receptive field in response to horizontal bar motion at all (Fig. 3d, third panel).

We next analyzed whether the preferred movement direction was correlated to morphological features of the recorded neurons. Neurons from group 1 were recorded most frequently and were hence investigated in most detail. For two cell types (LO-PN-bilat-1 and LO-PN-contra-1), all individuals receiving input in the left optic lobe preferred clockwise bar movements (Fig. 3c, top left, d, bottom), whereas individuals receiving input in the right optic lobe preferred counter-clockwise motion (only measured in LO-PN-bilat-1, Fig. 3c, top right). Interestingly, the third member of group 1 neurons (LO-PN-ipsi-1) showed the opposite behavior. It preferred counter-clockwise motion when receiving input in the left optic lobe, but preferred clockwise motion when receiving input on the right side (Fig. 3c, bottom). Consistent with this difference, both sets of neurons received input from different layers of the lobula, layer 5 for LO-PN-bilat-1 and -contra-1, in contrast to inputs focused on mostly layer 4 for LO-PN-ipsi-1. While consistent in group 1 neurons, group 2 neurons did not follow this rule. LO-PN-bilat-3 innervated layer 5 of the lobula, but showed a direction preference similar to LO-PN-ipsi-1 (a layer 4 neuron), and LO-PN-bilat-4 innervated layer 4 but did not have a pronounced horizontal motion tuning at all (Fig. 3d).

As some LO-PNs responded only weakly to horizontal bar motion, we tested whether these cells might be tuned to vertical motion instead (Fig.3e-g). Indeed, the two cell types that showed the weakest responses to horizontal motion exhibited strong, direction-selective responses to a moving horizontal stripe (LO-PN-bilat-2 and -bilat-4, Fig. 3g). While one preferred upwards movement (bilat-4), the other one preferred downwards movement (bilat-2). Interestingly, LO-PN-bilat-1 cells, which were highly sensitive to horizontal motion, were also excited by upwards motion and inhibited by downwards motion (Fig. 3f,g). This suggests that the true tuning direction of these cells is a combination of horizontal and vertical movements, such as those encountered during specific flight maneuvers.

After testing responses to localized movements of narrow bars, we next investigated the responses of LO-PNs to wide-field optic flow stimuli (Fig. 4). Generally, all neurons showed clear responses to wide-field motion and these responses were highly consistent with the described responses to a single bar. This means that wide-field motion responses could be predicted by the local motion tuning and receptive field size of the same cells.

The grouping of LO-PNs into a strongly responding and a weakly responding set of cell types was also supported by optic flow responses, albeit, LO-PN-bilat-2 could not be tested with these stimuli. Most group 1 cells (LO-PN-bilat-1, -ipsi-1, -contra-1) showed very strong responses with sustained peak activities of at least 100 impulses per second (Fig. 4a-c,e,f), while the two members of group 2 showed much weaker responses (Fig. 4g,h). Responses to yaw rotational optic flow in the preferred motion direction revealed increasing excitation strength with increasing speed of the optic flow pattern in all group 1 cells and in the LO-PN-bilat-3 (Fig. 4a-g). In line with the lack of responses to the vertical bar in the LO-PN-bilat-4 neuron, no responses to horizontally moving gratings were observed in this cell (Fig. 4h). For all other neurons, the stimulus velocity that led to maximal excitation was consistently ca. 100°/s, above which the firing rates of the cells did not increase further.

As all tuning curves were all obtained with a grating of identical spatial frequency, we next tested whether the observed dependence on grating velocity was due to speed tuning or temporal frequency tuning. To this aim we presented additional stimuli to a subset of neurons, in which we varied the spatial frequency at constant grating velocity (Supplemental Fig.1). For neurons tuned to temporal frequencies, a clear tuning to particular spatial frequencies would be expected, while speed tuned neurons should show constant responses to speed, independent of spatial frequency. Whereas the difference between simulated clockwise and counter-clockwise yaw rotations was preserved, we did not observe clearly peaked tuning curves in response to different spatial frequencies. Nevertheless, a trend towards lower firing rates at spatial frequencies above 0.1 cycles/° was visible in most cells. Although our data are more consistent with the hypothesis that the recorded neurons encode stimulus speed, more systematic testing of spatial frequency tunings at different speeds would be needed to completely rule out temporal frequency tuning.

To further characterize the optic flow responses of the LO-PNs, we observed the dynamics of the motion response to our stimuli. During all stimulus presentations we first switched the grating on in stationary mode for 0.5 s, after which the grating moved at constant velocity for 3 s before it was switched off again (Fig. 4). Before and after the stimulus the display was dark. Stimulus onset caused a strong lights-on response in all LO-PNs, irrespective of the subsequent motion response (Fig. 4a,b,e-h). For all motion responses in group 1 neurons, there was no observable adaptation of the response over the course of the 3 stimulus (Fig. 4b,e,f). In contrast, underlining their generally more variable responses, both group 2 neurons tested with optic flow stimuli showed different responses. LO-PN-bilat-3 was the only cell in which the strength of the response diminished over the course of the stimulus, indicating motion adaptation (Fig.4g). Finally, LO-PN-bilat-4, although not responding to horizontal optic flow, was the only cell with a strong lights-off response (Fig.4h). To generate these different response dynamics, both group 2 neurons likely receive qualitatively different input from upstream neurons compared to group 1 neurons.

### Medulla projection neurons

#### Morphology

Besides neurons connecting the lobula to the central brain, we also recorded from medulla projection neurons (ME-ME-PNs). These neurons projected from a specific layer of one medulla to the posterior protocerebrum and further into the contralateral medulla (Fig. 5). For all cells in which the soma was labeled, it was located near the optic stalk. The main neurite crossed the midline in the posterior optic commissure. All central brain arborizations of ME-ME-PNs were heavily blebbed (Fig. 5g), while medulla arborizations were fine ipsilaterally to the soma and blebbed on the contralateral side (Fig. 5c,d and e,f, respectively). These cells thus had a pronounced morphological polarity with likely input fibers in the ipsilateral medulla (near the soma) and outputs in the central brain and the contralateral medulla.

**Figure 5:**
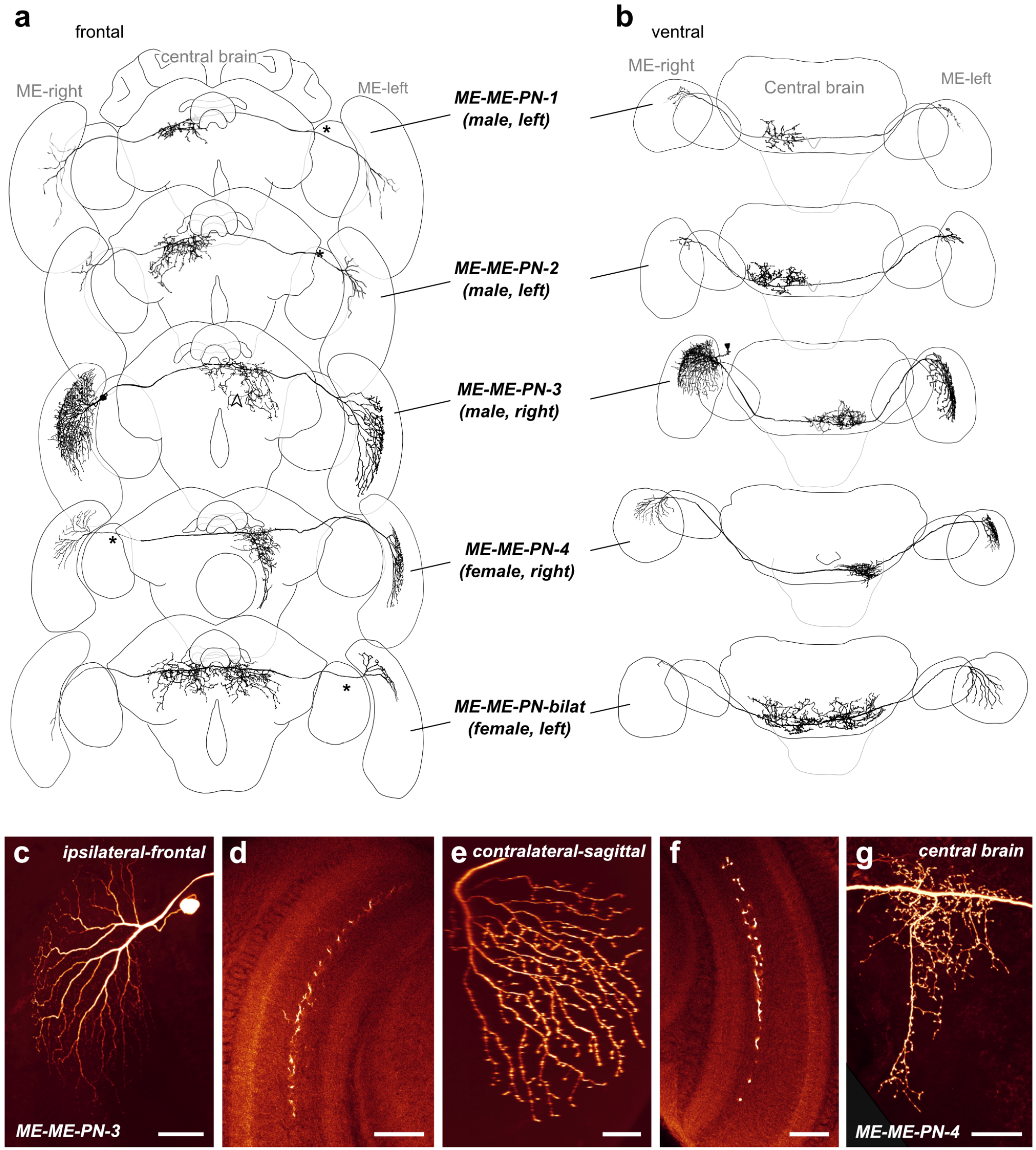
Morphology of large-field inter-medulla neurons with central brain projections (ME-ME-PNs). (a) 3D reconstructions of different ME-ME-PN types (frontal view), with schematic outline of the brain for orientation. Asterisks indicate approximate soma position. (b) Ventral view. (c-f) Confocal images of a neurobiotin-injected ME-ME-PN-3 neuron. (c) Maximal intensity projection of the cell body and arborizations in the ipsilateral medulla (frontal view). (d) Single optical section showing arborizations in a single layer of the ipsilateral medulla. (e) Maximal intensity projection of the arborizations in the contralateral medulla (sagittal view). (f) Single optical section showing arborizations in a single layer of the contralateral medulla. (g) Maximal intensity projection of the arborizations of a neurobiotin-injected ME-ME-PN-4 neuron in the posterior protocerebrum. Scale bars = 100 *µ*m (c), 50 *µ*m (d-g)

Generally, all arborizations in the medulla were concentrated in layer 6 of the outer medulla, but fibers occasionally protruded into the neighboring layer 5 and 7 as well (Fig.5d,f). While all types of ME-ME-PN neurons appeared similar in overall morphology, they differed in the details of their projection fields in the medulla and, more strongly, in the extent of the projection fields in the posterior protocerebrum. Using the latter feature, we divided all neurons into five types (Fig. 5a,b). Most different from the other types was the ME-ME-PN-bilat neuron, as it projected to large, bilateral regions in the protocerebrum. Three types (ME-ME-PN-1, ME-ME-PN-2, and ME-ME-PN-4) projected exclusively contralaterally, whereas the remaining type (ME-ME-PN-3) projected predominantly contralaterally but had additional fibers around the brain midline.

In more detail, the smallest projection fields were found in the ME-ME-PN-1 cell (n=1), focused in the posterior part of the ventromedial protocerebrum. Slightly larger projection fields, more substantially extending into the posterior ventrolateral protocerebrum, defined the ME-ME-PN-2 cells (n=2). Distinct from the two latter cell types, ME-ME-PN-3 cells (n=2) innervated the posterior inferior and ventromedial protocerebrum. Whereas arborizations of ME-ME-PN-1 and -2 neurons extended anteriorly with some arborizations at the level of the CX noduli, ME-ME-PN-3 cells lacked these fibers but possessed characteristic arborizations posterior to the CX, bilaterally arranged around the brain midline. The most frequently encountered cell type was the ME-ME-PN-4 neuron (n=8), which shared output projection fields in the ventromedial protocerebrum with the remaining types, but was characterized by additional fibers projecting further ventrally into contralateral, peri-esophageal neuropils (Fig. 5a,g). Finally, the ME-ME-PN-bilat neuron (n=2) was the only cell type among medulla projection neurons with bilaterally symmetrical central projections. On both sides of the midline, these projections covered posterior regions of the ventrolateral and ventromedial protocerebrum.

Near the soma, the main neurite of all ME-ME-PN cells branched into a fan of arborizations that penetrated the ipsilateral medulla. On this input side, the projections in the innervated medulla layer were generally focused on the dorsal half of the neuropil, but extended beyond the equator for ME-ME-PN-3 cells (Fig.5). ME-ME-PN-1 cells was the only cell type with inputs restricted to a ventral region. As on the input side, fibers were restricted to a single medulla layer on the output side (layer 6 with some excursions into neighboring layers). Owing to faint staining, these arborizations could not always be traced completely (Fig.5, ME-ME-PN-2, ME-ME-PN-bilat), leaving some uncertainty about the full extent of the projection fields. The center of the radiating point of the contralateral medulla branches nevertheless indicated the focus of those fibers. While outputs were generally located in a dorso-ventral position that reflected the position of the corresponding fibers on the input side, a shift towards more ventral regions was also regularly observed. The extent of this shift ranged from subtle in ME-ME-PN-4 neurons to clearly medial regions in ME-ME-PN-3 cells. It was extreme in the bilateral ME-ME-PN-bilat cell type, in which the outputs were shifted to the opposite side the medulla compared to the dorsally located inputs.

#### Physiology

The recordings from medulla projection neurons yielded physiological characteristics that were distinct from those of lobula projection neurons in several key aspects. To characterize these cells, we tested the same stimuli as described for the lobula neurons. Mapping of the receptive field with a bright vertical bar was carried out in all recordings, and responses to a range of optic flow stimuli were tested in most cell types (Fig. 6 and Suppl. Fig. 2,3).

**Figure 6:**
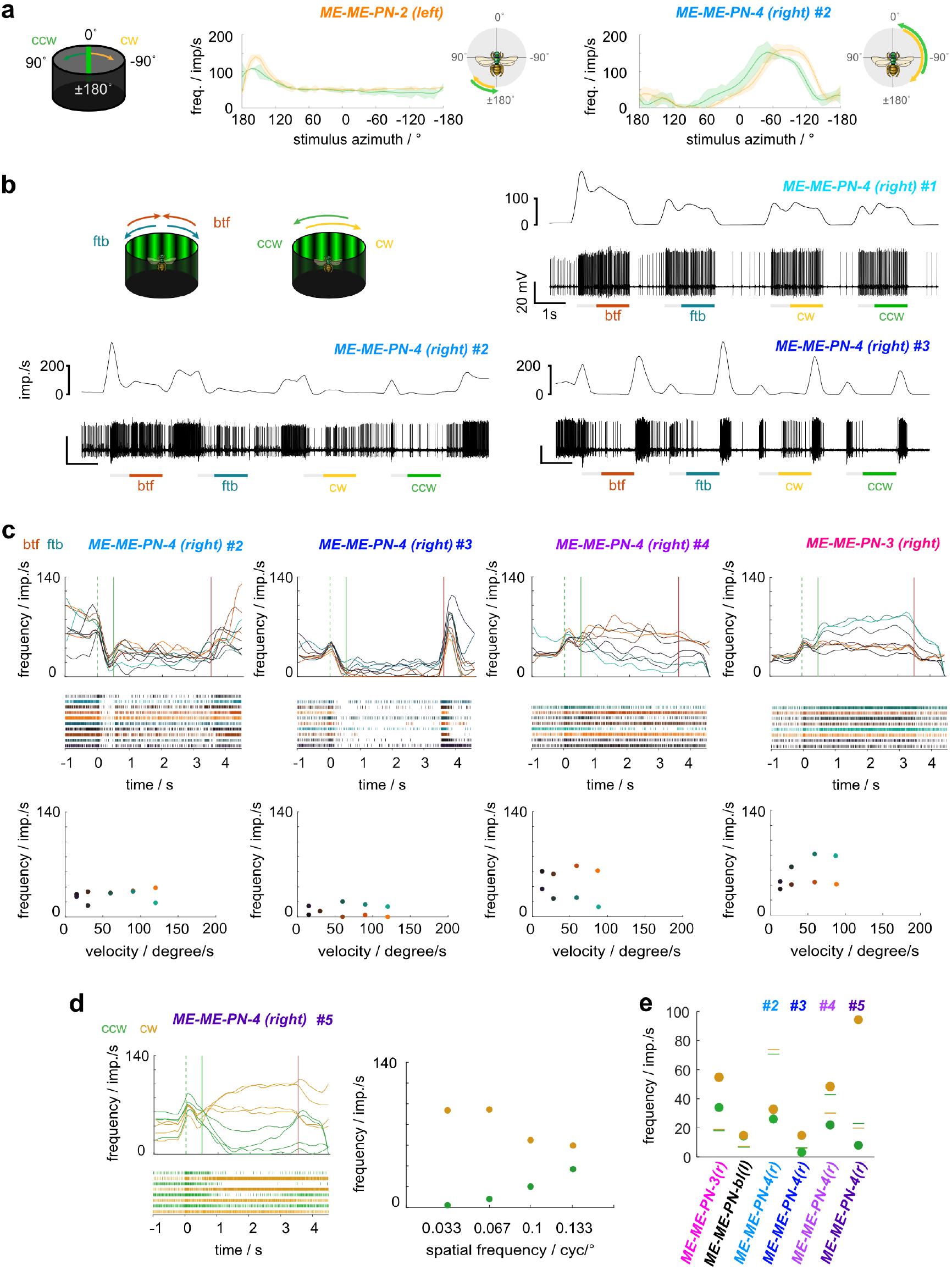
Physiology of tangential inter-medulla neurons (ME-ME-PNs). (a) Normalized average response curves of a ME-ME-PN-2 and a ME-ME-PN-4 neuron of the right hemisphere during receptive field mapping with a green vertical stripe moving clockwise (cw, yellow) or counter-clockwise (ccw, green). Cartoons on the right: approximate horizontal extent of each receptive field and preferred movement directions. (b) Example responses of three individual ME-ME-PN-4 neurons (#1-#3) to front-to-back (ftb) and back-to-front (btf) optic flow and cw and ccw optic flow. Bottom: spike train; top: sliding average; gray bars below spike trains: stationary gratings; colored bars: gratings moving at constant speed. (c) Top: Activity of individual ME-ME-PN-4 in response to ftb and btf optic flow of different velocities (dark to light cyan/orange; for speed values see bottom graphs). Green dotted lines: grating presented; green solid lines: motion onset; red lines: motion stop. Middle: Corresponding raster plots. Bottom: Mean frequency during the final 2 s of each stimulus bout; color of dots as in top graphs. (d) Response of an individual ME-ME-PN-4 neuron (#5) to cw (yellow) and ccw (green) rotational optic flow of different spatial frequencies. Right: mean frequency during the final 2 s of each stimulus bout. (e) Mean frequency of individual ME-ME-PNs during the final 2 s of cw (yellow) and ccw (green) rotational optic flow with a spatial frequency of 0.067 cyc/°. Green and yellow lines indicate mean background activity during 2 s preceding each stimulus. No ND filter in (b, #3; c, #3,#4, and ME-ME-PN-3; d; e, ME-ME-PN-3, #3-#5)

In all recordings, responses to a moving vertical bar had a dominant excitatory component and, importantly, were independent of the movement direction (Fig. 6a, Suppl. Fig. 2,3). In the majority of recordings, the response peaks were offset between clockwise and counter-clockwise movements of the bar. As this shift occurred in the direction of movement, it is consistent with being caused by delays in signal processing between the photoreceptors and the recorded medulla cells (Fig.6a). The receptive fields mapped in this way were always concentrated on the side of the brain containing the neuron’s soma. In most neurons, the receptive fields were adjacent to the midline and extended slightly into the contralateral field of view (Fig. 6a, right). Exceptions to this were the ME-ME-PN-2 (Fig. 6a, left) and the ME-ME-PN-3 cells (Suppl. Fig. 3d,e). In both ME-ME-PN-2 neurons the receptive fields were located in the posterior field of view, with the smallest width of all medulla PNs (ca. 60°) (Fig.6a, Suppl. Fig. 2). The receptive fields of both ME-ME-PN-3 neurons were shifted away from the anterior midline towards posterior directions, but to different degrees. One mirrored the shape and location of receptive fields of ME-ME-PN-2 neurons (Suppl. Fig. 3d), while the other one responded in a wider range in the ipsilateral field of view (Suppl. Fig. 3e). This inconsistency between the anatomical identity and the physiological features of the neurons was also visible within the seven physiologically characterized ME-ME-PN-4 cells. Here, the largest receptive fields spanned approximately 180°, while the narrowest example was only around 80° wide (Fig6. b, Suppl. Fig. 2). Furthermore, three of these neurons had more complex transient lights-on responses, in which a bout of inhibition followed the initial excitatory burst. In these neurons a strong contralateral inhibition was additionally observed during receptive field mapping with a bright bar, potentially a result of an inhibitory rebound from the preceding strong ipsilateral excitation (Suppl. Fig. 2d,f,g).

In four recordings, the vertical receptive field extent was probed with a moving horizontal stripe. It elicited an excitatory response in all cases, of which three were independent of movement direction (ME-ME-PN-bilat cells and the ME-ME-PN-1 cell, Suppl. Fig. 3c,f,g). Surprisingly, one of the ME-ME-PN-4 cells responded only to upwards movements with an excitation, but was slightly inhibited by downwards movement (Suppl. Fig. 3f).

The responses to optic flow stimuli were remarkably inconsistent both between neuron types and within types (Fig. 6, Suppl. Fig. 2,3), suggesting that the anatomically defined cell types either comprise multiple distinct functional subtypes, or that the responses vary due to parameters not under experimental control. Generally, responses to wide-field motion were weaker than the responses to the vertical bars when tested in the same recordings. While most recordings revealed excitatory responses to wide-field optic flow stimuli (Fig. 6b, top right,c, right), consistent with excitatory responses to bar stimuli, the responses were inhibitory in two cases (Fig. 6b (bottom), c (cells #2, #3)). Interestingly, this flip in response sign was observed between recordings of the same cell type (ME-ME-PN-4, Fig.6b). Further substantiating the variable response features, we found direction-selective responses to optic flow stimuli in three recordings (Fig. 6c, cell #4 and ME-ME-PN-3,d), despite the fact that moving bars always elicited excitation in both directions during horizontal movement. Finally, a consistent characteristic of all neurons was a strong lights-on excitation, which was often most pronounced during the first stimulus presentation (Fig.6b). Yet, even these phasic properties were variable, as some neurons showed a pronounced inhibition following the onset excitation, and one neuron possessed a strong lights-off excitation in addition to the consistent lights-on response (Fig. 6b and c (cell #3)).

Given that ME-ME-PN cells interconnect both medullae, it was conceivable that contralateral stimulation could affect the ipsilateral optic flow responses, leading to potentially different responses when simulating yaw rotations or translational movement, despite the fact that both stimuli would appear identical to the unilaterally restricted receptive fields of these cells. However, in our initial tests at the beginning of each recording, no qualitative difference was found in response to both stimuli with the same underlying spatial frequency and movement speed (Fig.6b). As we tested progressive and regressive velocity series more frequently in these neurons than velocity series of yaw rotations, we used this stimulus to describe the responses of ME-ME-PNs to different grating velocities. Based on these data, only one ME-ME-PN-3 neuron showed an increase in response strength with stimulus speed, with front-to-back optic flow leading to a stronger response than back-to-front optic flow, saturating already at 60° per second (Fig. 6c, ME-ME-PN-3). While a difference between front-to-back and back-to-front optic flow was observed in three out of nine recordings, the response profiles did not show speed dependency (Fig. 6c (cells #3, #4); Suppl. Fig. 2a,b,c). For the neurons in which clockwise and counter-clockwise yaw rotations were tested at different speeds (3 recordings), data were identical to progressive/regressive optic flow velocity data (Suppl. Fig. 2a,b, Suppl. Fig. 3e).

As we found very few responses to different optic flow velocities, we additionally tested the responses of ME-ME-PN cells to different spatial frequencies of optic flow presented at 60° per second, both for clockwise and counter-clockwise yaw rotations (Fig. 6d, Suppl. Fig. 2,3). Except for one ME-ME-PN-4 cell (#5; Fig.6d), no cell changed their overall firing frequency in response to different spatial frequencies (Supplemental Fig.2,3). However, these tests confirmed the surprising direction selectivity of optic flow responses identified with front-to-back and back-to-front optic flow (Fig.6e).

Finally, in contrast to lobula projection cells, motion adaptation was found in almost half the recorded neurons during progressive and regressive optic flow stimulation (four out of nine recordings) (e.g. Fig. 6d).

### Inter medulla/lobula neurons

In addition to projection neurons from the lobula and medulla, we found one cell type that interconnected the medulla and the lobula of both hemispheres (MELO-MELO-PN, n=3;Fig.7), which likely corresponds to the serpentine neuron described by Hertel and Maronde (1987). The soma of each neuron was located near the optic stalk in the anterior protocerebrum. The primary neurite projected toward the ipsilateral lobula and split into several branches. One branch projected via the superior optic commissure into the central brain, while the other branches gave rise to fine, input-like arborizations around layer 2 of the lobula (Fig.7b,c, arrows). The projection in the central brain split up again, leading to one branch that continued contralaterally and one branch that projected back into the ipsilateral optic lobe. The latter gave rise to blebbed processes throughout the ventral half of the outer lobula and the outermost layer of the inner medulla (layer 8, Fig 7b,c). The contralateral projections covered the same regions in the contralateral medulla and lobula with equally blebbed ramifications (Fig.7d). Additionally, all three neurons possessed sparse, blebbed arborizations that extended either only contralaterally (n=1) or bilaterally (n=2) from the main neurite into several regions of the central brain, including inferior and ventrolateral neuropils.

**Figure 7:**
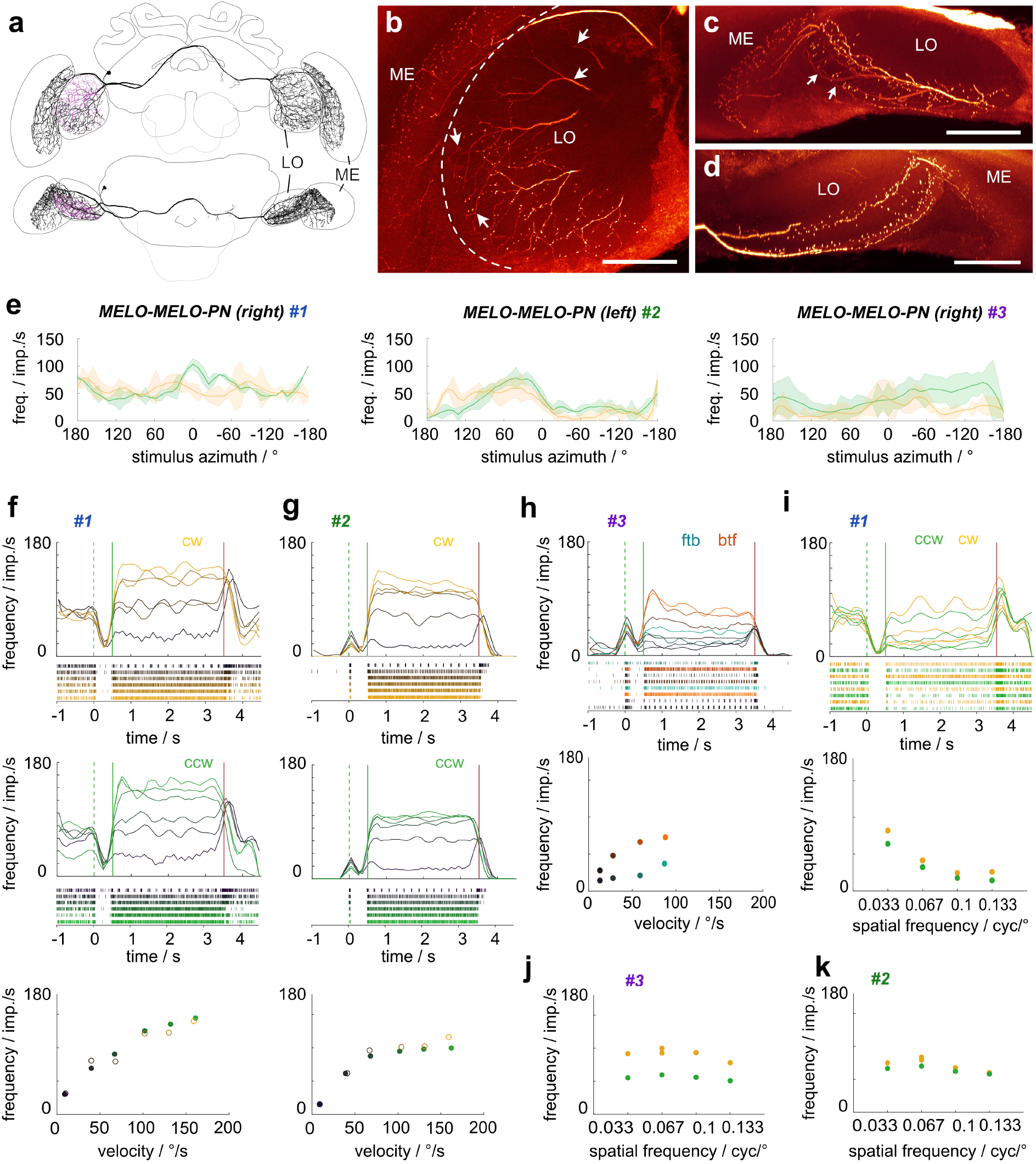
Anatomy and physiology of inter-medulla/lobula neurons (MELO-MELO-PNs). (a) Frontal (top) and ventral (bottom) view of the 3D reconstruction of a MELO-MELO-PN. (b-d) Maximal intensity projection of a few optical sections of the confocal stack illustrating the arborizations of a neurobiotin-injected MELO-MELO-PN in the ipsilateral lobula and medulla (b,c) and the contralateral lobula and medulla (d). Frontal view (b), ventral view (c,d). Arrows indicate smooth fibers. (e) Average response curves of three individual cells during receptive field mapping. Yellow: cw movement, green: ccw movement. (f) Activity of the MELO-MELO-PN #1 of the right hemisphere in response to cw (yellow) and ccw (green) wide-field optic flow of different velocities (dark to light yellow/green, values: 10°/s, 40°/s, 70°/s, 100°/s, 130°/s, and 160°/s). Green dotted lines: grating presented; green solid lines: motion onset; red lines: motion stop. Bottom graph: mean response frequency during the final 2 s of each stimulus bout of the cw and ccw stimulus sequence. (g) Same as in (f) but for the MELO-MELO-PN #2 of the left hemisphere. (h) Activity of the MELO-MELO-PN #3 of the right hemisphere in response to front-to-back (ftb) and back-to-front (btf) optic flow of different velocities (dark to light cyan/orange, values: 15°/s, 30°/s, 60°/s, and 90°/s). (i) Activity of the MELO-MELO-PN #1 in response to cw (yellow) and ccw (green) optic flow of different spatial frequencies. Bottom graph: mean response frequency during the final 2 s of each stimulus bout. (j,k) Same as in the bottom graph in (i) but for the other two MELO-MELO-PNs. All stimuli without ND filters.

The three MELO-MELO-PN cells were tested with the same stimuli as the previously described neurons and showed generally very strong motion responses (Fig. 7e-k). Receptive field mapping with a vertical bar moving around the animal revealed localized receptive fields in the ipsilateral field of view. These were much noisier than for the lobula and medulla projection neurons. While they were all located adjacent to the midline, they extended posteriorly to different degrees (Fig.7e). Two of the cells showed weak direction selectivity (Fig. 7e, cells #1,#3), while the third one was equally excited by both movement directions (Fig. 7e, cell #2).

In contrast to the poorly defined receptive fields, the optic flow responses were strong and clear (Fig.7f-i). All cells showed a pronounced response to different stimulus velocities, either in response to clockwise and counter-clockwise yaw rotations (tested in two cells, Fig. 7f,g) or progressive and regressive optic flow (tested in one cell, Fig. 7h). Yaw rotational optic flow responses were not direction-selective, non-adapting and saturated at similar levels as the LO-PNs (around 100 ° /s). The cell tested with progressive and regressive optic flow approached its saturation plateau between 60 and 100 ° /s, was weakly direction-selective, and showed signs of motion adaptation at the highest velocities (Fig.7h). The neurons differed in their background activity, which was high (around 60-80 imp/s) in one cell (Fig. 7e, #1), but low in the others (below 10 imp/s; Fig.7e, #2, #3). Interestingly, this difference did not affect the absolute response levels to the stimuli, leading to an apparent inhibition in the cell with high background activity (for low stimulus velocities) and only excitatory responses in the other two cells. All cells were also tested with different spatial frequencies, leading to no clear tunings, but yielding a consistent trend towards preference of lower spatial frequencies (Fig. 7i-k).

Different from LO-PNs, the MELO-MELO-PN cells showed a complex phasic lights-on and lights-off response. Stimulus onset consistently caused an excitatory burst, followed by a near complete inhibition until motion onset (Fig. 7f-i). The end of the motion stimulus (lights-off) caused an excitatory burst at low velocities, which became inhibitory at higher velocities. Note that the inhibitory component was hidden in the neurons with low background activity (Fig. 7g,h), while the excitatory onset was hidden in the cell with high background activity (Fig.7f). Consistent with these complex phasic responses, the lowest grating velocity resulted in pronounced, regular activity bursts during stimulus display, suggesting entrainment of the neuron to individual stripes of the grating. Entrainment to stripes was also observed with the spatial frequency stimuli (Fig.7i). This response pattern was generally not observed in lobula projection neurons, indicating that the neural mechanisms of constructing the motion responses in both sets of neurons is likely distinct. This idea is in line with the fact that both sets of cells receive their input in different layers of the lobula (layer 2 in MELO-MELO-PN cells and layers 4 or 5 in most LO-PNs) and therefore from different upstream neurons.

### Centrifugal feedback neurons

Although our study focused on neurons that provide information from the optic lobes to the central brain, we obtained several recordings from neurons with the opposite polarity, i.e. suggesting information flow from the central brain back into the optic lobes. These centrifugal neurons likely play a role in feedback and possessed very different response properties compared to the projection neurons. Overall, we found two cell types that project from the posterior protocerebrum back to the lobula (LO-CNs, Fig. 8) and four that project from similar regions of the central brain to the medulla (ME-CNs, Fig. 9). With the exception of ME-CN-2 neurons (n=2), each cell type was encountered only once.

**Figure 8:**
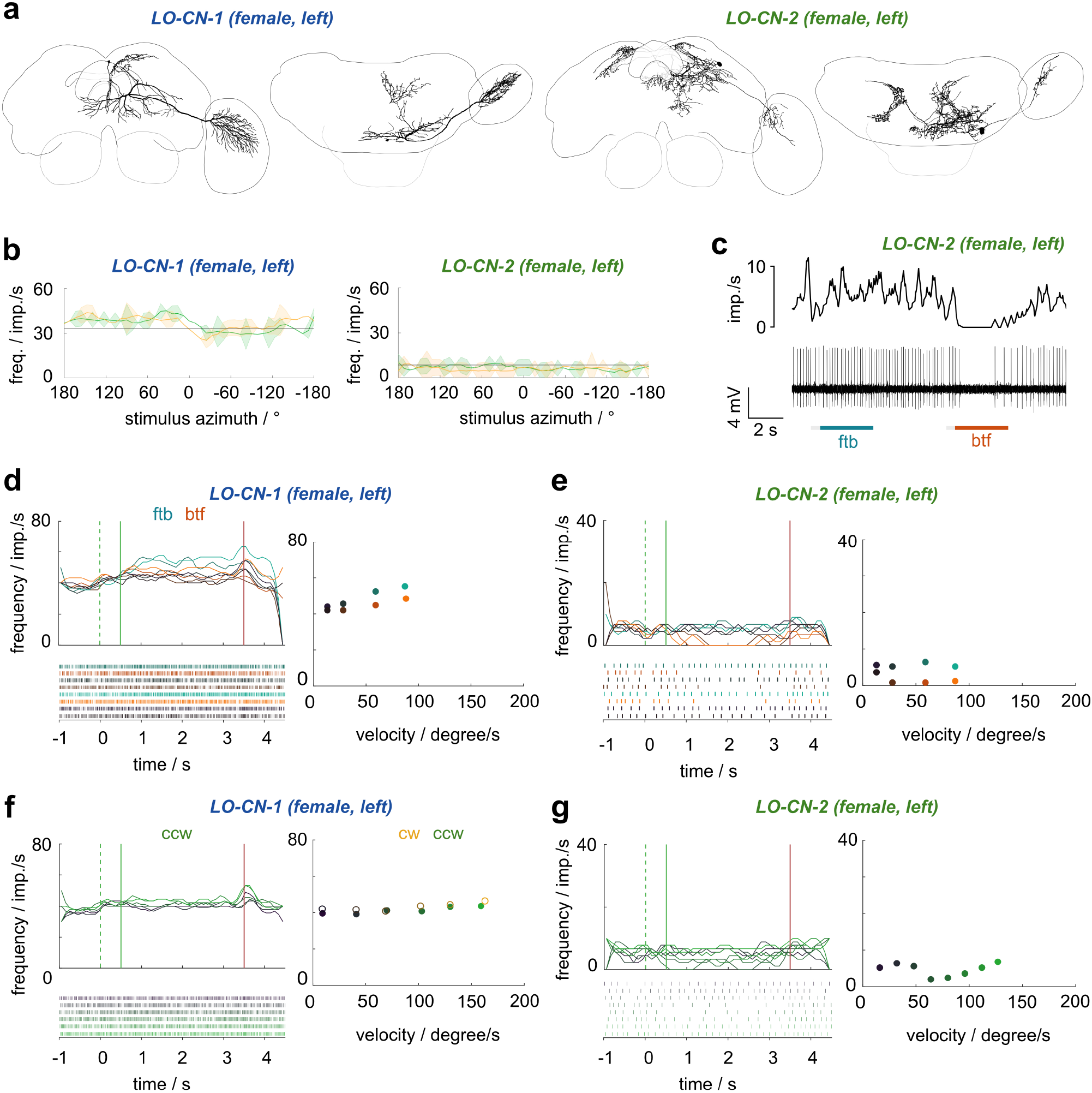
Anatomy and physiology of centrifugal feedback neurons with arborizations in the lobula (LO-CNs). (a) 3D reconstructions of different LO-CN types with schematic outline of the brain for orientation. Left images: frontal view; right images: ventral view. (b) Average response curves of the LO-CN cells during receptive field mapping. Yellow traces: clockwise (cw) movement; green traces: counter-clockwise (ccw) movement; gray horizontal line: mean background activity during 2 s preceding the stimulus. (c) Example response of a LO-PN-2 neuron in response to front-to-back (ftb) and back-to-front (btf) optic flow. Bottom: spike train; top: sliding average; gray bars below spike trains: stationary gratings; colored bars: gratings moving at constant speed. (d) Activity of the LO-CN-1 neuron in response to ftb and btf optic flow of different velocities (dark to light cyan/orange; for speed values see graphs on the right). Green dotted lines: grating presented; green solid lines: motion onset; red lines: motion stop. Graphs on the right: Mean frequency during the final 2 s of each stimulus bout; color of dots as in graphs on the left. (e) Same as in (d) but for the LO-CN-2 neuron. (f) Activity of the LO-CN-1 cell in response to clockwise (yellow) and counter-clockwise (green) optic flow of different velocities (dark to light yellow/green; for speed values see graph on the right). Graph on the right: mean frequency during the final 2 s of each stimulus bout; color of dots as in graphs on the left. No ND filter in (b-f).

**Figure 9:**
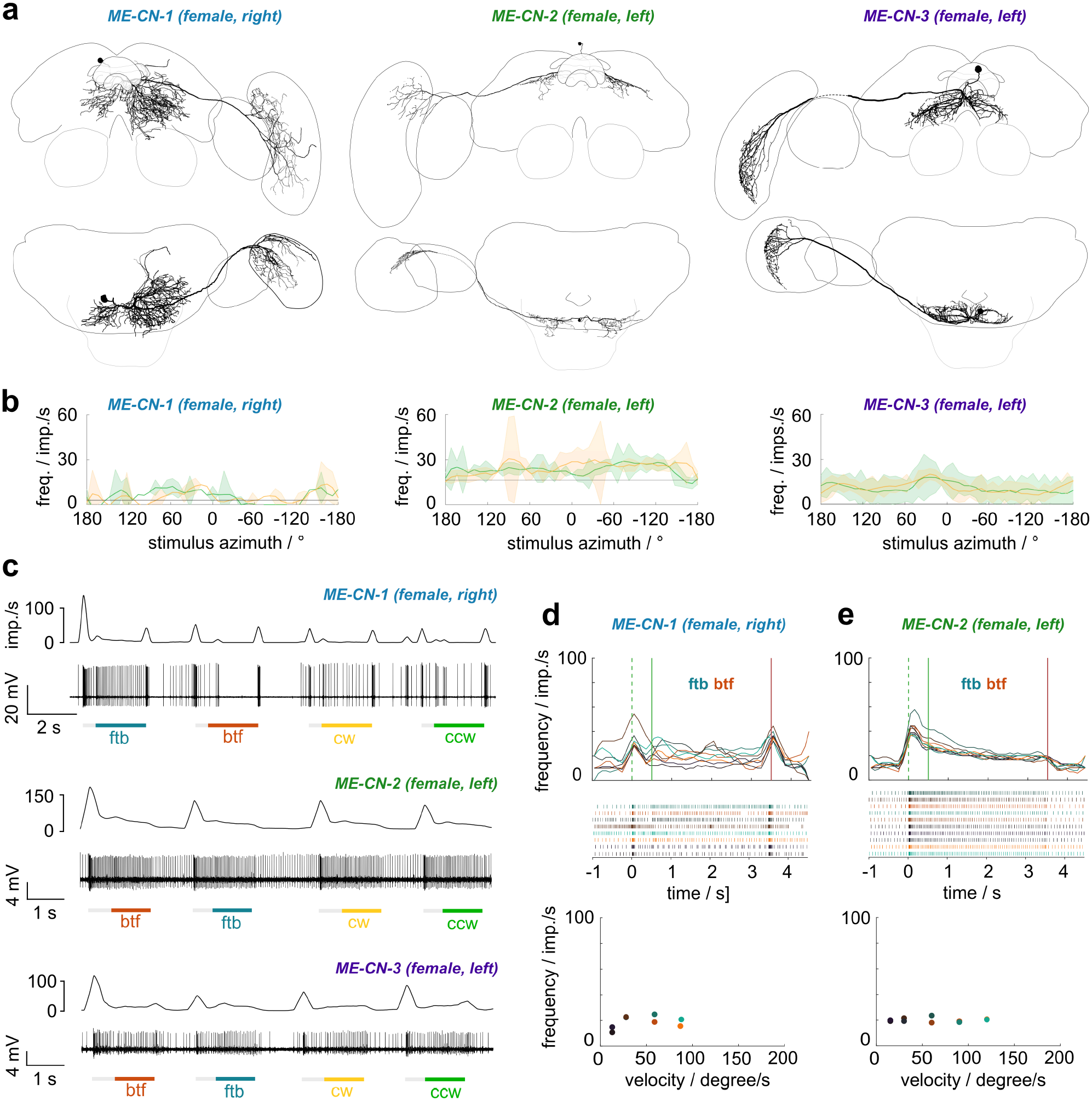
Anatomy and physiology of centrifugal feedback neurons with arborizations in the medulla (ME-CNs). (a) 3D reconstructions of different ME-CN types with schematic outline of the brain for orientation. Top row: frontal view; bottom row: ventral view. (b) Average response curves of the ME-CN cells during receptive field mapping. Yellow traces: clockwise (cw) movement; green traces: counter-clockwise (ccw) movement; gray horizontal line: mean background activity during 2 s preceding the stimulus. (c) Example responses of the neurons in response to front-to-back (ftb) and back-to-front (btf) optic flow and cw and ccw wide-field optic flow. Bottom: spike train; top: sliding average; gray bars below spike trains: stationary gratings; colored bars: gratings moving at constant speed. (d) Top: Activity of the ME-CN-1 and a ME-CN-2 neuron in response to ftb and btf optic flow of different velocities (dark to light cyan/orange; for speed values see bottom graphs). Green dotted lines: grating presented; green solid lines: motion onset; red lines: motion stop. Middle: Corresponding raster plots. Bottom: Mean frequency during the final 2 s of each stimulus bout; color of dots as in top graphs. No ND filter for ME-CN-1 in (b-d).

Cell bodies of both LO-CN neurons (Fig. 8a) were located dorsally in the posterior protocerebrum of the left hemisphere. From the cell body the primary neurite of the PN-LO-1 neuron projected ventrally and gave rise to several branches that formed arborizations mainly in the ipsilateral inferior, ventromedial and ventrolateral protocerebrum, with some processes extending into the contralateral inferior protocerebrum. Additionally, one branch projected dorso-anteriorly and gave rise to arborizations in the ipsilateral superior protocerebrum. The primary neurite of the PN-LO-2 neuron ran medially and gave rise to arborizations that innervated the ipsilateral inferior, ventromedial and ventrolateral protocerebrum. Two branches ran dorsally and arborized bilaterally in the ipsi- and contralateral superior protocerebrum and also provided arborizations to the contralateral inferior protocerebrum.

In the optic lobe, both cells had varicose arborizations in dorsal regions of the inner lobula, distinct from the input fibers of the LO-PNs in the outer lobula. The projection field of the LO-CN-2 neuron within the lobula was located within layer 8 and was relatively small compared to the arborization area of the LO-CN-1 neuron, which extended from proximal to distal layers of the inner lobula.

Responses to all presented motion stimuli were subtle in both LO-CN cells (Fig.8b-g). Neither neuron had a clear receptive field when tested with a vertically moving bar. In the LO-CN-1 neuron, a slight direction-independent excitation was present across the entire left visual field, consistent with the brain hemisphere in which the neuron arborized (Fig. 8b, left). In the LO-CN-2 neuron the same stimulus led to no obvious response (Fig. 8b, right).

In both neurons, responses to wide-field stimuli were also much weaker than in LO-PNs and no obvious relation was evident between the bar stimulus and the properties of the optic flow responses. Yaw rotational optic flow only very slightly raised the mean firing rate above background level of one of the neurons (Fig.8f), while it caused comparably strong inhibition for counter-clockwise stimulations in LO-CN-2, but only at medium grating velocities (Fig.8g). Interestingly, front-to-back and back-to-front optic flow was more consistently effective in driving these neurons. The LO-CN-2 neuron was specifically inhibited by back-to-front optic flow, but did not respond to front-to-back optic flow, in line with the directional response to simulated yaw rotations and with input on the right side of the brain, opposite to the lobula branches (Fig. 8c,e). In the LO-CN-1 neuron, both, progressive and regressive optic flow led to a speed-dependent excitation during the motion stimulus. This response was stronger during front-to-back optic flow (Fig. 8d).

Response dynamics were also distinct from projection neurons and we never observed transient lights-on or lights-off responses, suggesting that these neurons are further removed from the sensory periphery. This and the observed responses to progressive and regressive optic flow is consistent with a possible role of LO-CN neurons in reporting higher level information, such as flight status, to neurons of the optic lobe.

Beside lobula centrifugal neurons, we also obtained recordings from three types of medulla centrifugal neurons: ME-CN-1, ME-CN-2 (n=2), and ME-CN-3 (Fig. 9a). Cell bodies of all three types were located in the posterior protocerebrum, superior to the PB and close to the brain midline. From the cell body the primary neurite of all cell types projected ventrally and split into two (ME-CN-2) or several (ME-CN-1, ME-CN-3) branches that gave rise to bilateral arborizations in the central brain. Arborizations in the medulla were predominantly blebbed, while central brain projections were characterized by fine arborizations intermingled with fewer blebbed processes.

The ME-CN-1 neuron formed numerous arborizations in the contralateral inferior, ventromedial, ventrolateral, and peri-esophageal neuropils, which extended anteriorly from the posterior brain border to the level of the CX, with some fibers reaching further into the lateral accessory lobes and the superior protocerebrum. In the ipsilateral hemisphere, arborizations were more restricted to areas close to the brain midline but still covered large parts of the posterior inferior and ventromedial protocerebrum. One prominent side branch originated ventrally to the PB and projected through the posterior optic commissure into the contralateral optic lobe. The neuron projected to a proximal region in the inner medulla and more sparsely innervated a distal layer of the outer medulla (likely layer 1). All fibers were limited to anterior portions of the medulla, but extended along the full dorsal-ventral axis. Arborizations of the ME-CN-2 neuron type were more restricted in the central brain and the optic lobe. The neuron innervated smaller areas in the inferior protocerebrum of both hemispheres along the ventral boundary of the posterior optic commissure. Only some processes extended into more ventral regions. Via the posterior optic commissure ME-CN-2 cells projected into the contralateral optic lobe, where anterior-dorsal regions of a medial layer of the medulla were innervated, possibly overlapping with ME-ME-PN arborizations. However, due to faint staining the layer identity could not be unambiguously determined. Finally, the ME-CN-3 cell had bilateral arborizations in the inferior and ventromedial protocerebrum as well as the peri-esophageal neuropils. In the contralateral hemisphere arborizations extended slightly further anteriorly and laterally than on the ipsilateral side. One prominent side branch ran via the posterior optic commissure toward ventral regions of the medulla, where fibers were present in layer 7. These fibers covered the full extent of the medulla along the anterio-posterior axis and thus provide direct overlap with the input and output fibers of the above described inter medulla projection neurons.

The physiological responses of ME-CN cells were different from LO-CN cells, but also much weaker in response strength compared to projection neurons (Fig.9b-e). In response to a moving vertical bar all three neuron types showed elevated activity in restricted azimuth ranges, but receptive field borders were blurred by noisy background activity (Fig.9b). In all cases the responses were not dependent on movement direction and were located adjacent to the anterior midline, extending between 60° and 120° posteriorly.

Responses of the PC-ME-1 neuron to wide-field stimuli were dominated by transient lights-on and lights-off responses, whereas the actual motion only elicited weak responses (Fig. 9c-e). While ME-CN-1 cells showed both on and off transients, the other two cell types only showed lights-on transients. These two cells also showed a weak excitation during movement, which was neither dependent on direction nor different between simulated yaw rotation and simulated translational movement (Fig. 9c,e). For the ME-CN-1 cell, responses were more variable and most pronounced during the first stimulus presentation (Fig.9c).

Overall, the more transient responses and the detectable receptive fields might indicate that the medulla centrifugal neurons receive more direct visual input compared to the lobula centrifugal cells. Together with the overlapping branches with medulla projection neurons both in the medulla and the central brain, these cells could thus form direct feedback loops to shape medulla motion responses.

### Overall projection patterns

Across all cell types we found evidence of three routes of information flow between the *Megalopta* optic lobe and the central brain. The first one (group 1 LO-PNs) was a strongly direction-selective pathway leading from layers 4 and 5 of the lobula to posterior regions of the protocerebrum, either ipsilaterally, contralaterally or bilaterally (Fig.10a). Responses of the three neuron types representing this pathway were highly stereotypical, with strong activity changes to both single bars and wide-field motion and defined receptive fields in the ipsilateral field of view. Similar to the input neurons to the noduli of the central complex (TN neurons, Stone et al., 2017), these cells saturated at grating velocities of around 100°/s.

**Figure 10:**
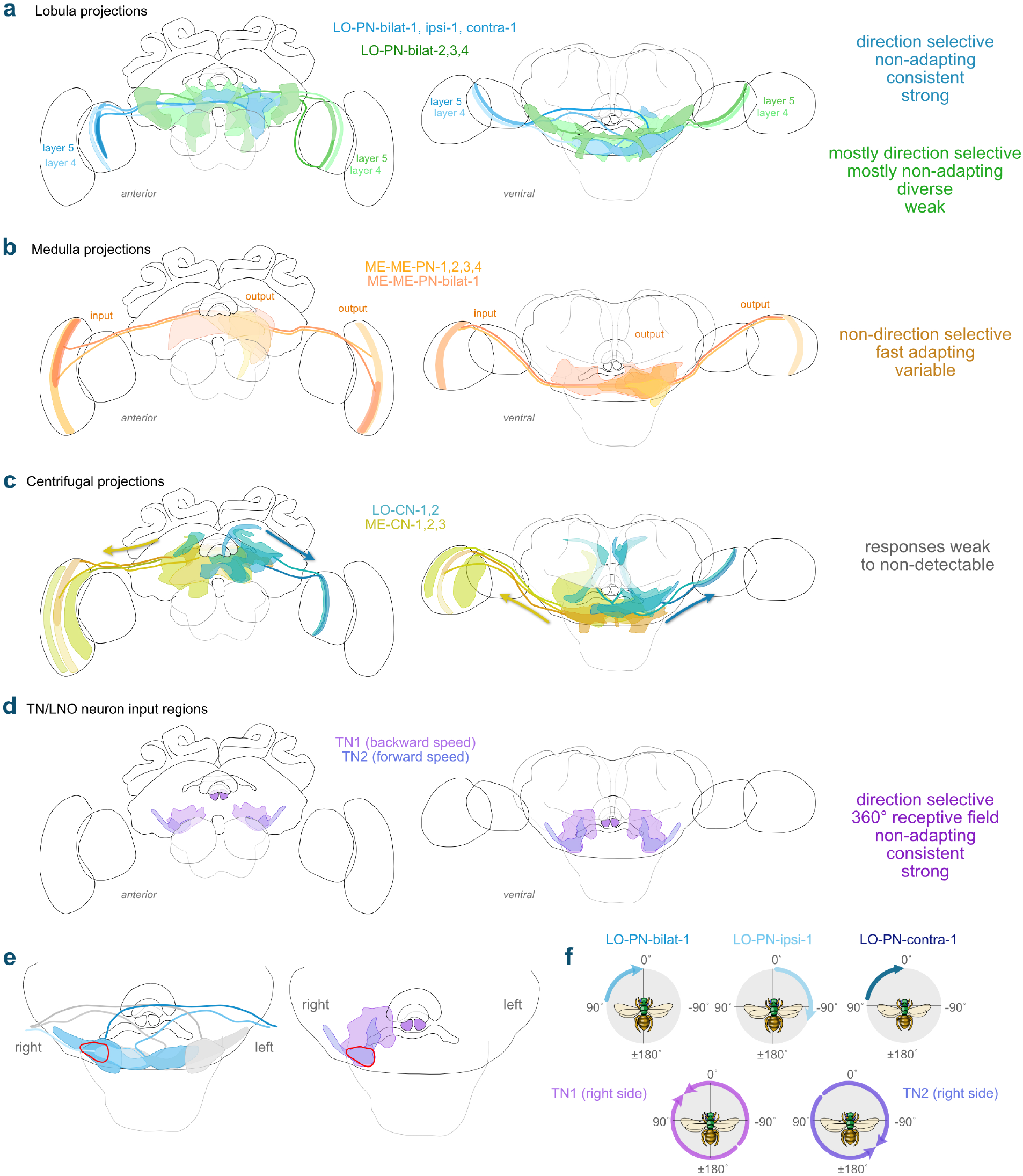
Major projection patterns of motion-sensitive neurons in *Megalopta*. (a) Two groups of lobula projection neurons (LO-PNs) with distinct physiological properties establish connections between layer four and five of the lobula and the central brain. Group 1 (shades of blue) LO-PNs (bilat-1, ipsi-1, and contra-1) share arborization areas in the posterior protocerebrum mostly omitted by arborizations of LO-PN-bilat-2, and -3 neurons (group 2, shades of green). The LO-PN-bilat-4 (group 2) innervates large areas of the posterior protocerebrum, overlapping with group 1 LO-PNs and the remaining group 2 LO-PNs. (b) Several types of medulla projection neurons connect layer six of the medulla to overlapping regions within the posterior protocerebrum. (c) Both, the lobula and the medulla receive feedback from wide areas within the posterior and medial protocerebrum via centrifugal medulla cells (ME-CNs, shades of turquoise) and centrifugal lobula cells (LO-CNs, shades of yellow). (d) Motion-sensitive central complex neurons (TNs) provide input from subregions of the posterior and medial protocerebrum to the noduli of the central complex (Stone et al., 2017). These neurons encode forward and backward translational movement velocity, mediated by translational optic flow. (e) Arborizations of group 1 LO-PNs and TN neurons partly overlap within the posterior protocerebrum (red ellipse). (f) The receptive field structures of group 1 LO-PNs together with TN neuron receptive fields. The combined input from LO-PNs cannot fully explain the complex responses of TN neurons (Stone et al., 2017).

The second pathway also originated in layers 4 and 5 of the lobula but showed weaker and more variable responses. The neurons contributing to this pathway either terminated in regions anteriorly of those from the first pathway as well as in regions surrounding the esophagus, or covered wide regions that included some of the projection fields of the first group (Fig.10a). While neurons of group 1 that originated in layers 4 and 5 of the lobula had opposite horizontal motion tunings, this was not the case for group 2 neurons from these layers, some of which were tuned to vertical motion or in the opposite direction to their group 1 counterparts from the same layer.

The third pathway comprised the medulla projection neurons, linking a single layer of the outer medulla to large regions of the posterior protocerebrum. While the projection fields overlapped with those of the lobula projection neurons, the encoded information was very different (Fig.10b). These neurons strongly responded to horizontal motion of a narrow bar, but in a not direction-selective manner. Responses to wide-field optic flow stimuli were remarkably variable and ranged from direction-selective sustained excitations to short transient inhibitions. Many medulla projection neurons showed clear signs of motion adaptation, a feature that was rarely observed in lobula projection neurons (and never in group 1 cells).

We found two additional pathways that transfer information between the optic lobes of both brain hemispheres rather than between the optic lobe and the central brain. Firstly, medulla projection neurons additionally interconnected both medullae, thereby sending a copy of the signals they relay from the ipsilateral medulla to the central brain also to the contralateral medulla (Fig. 10b). Secondly, a single neuron type (MELO-MELO-PN) linked layer 2 of one lobula to wider regions of the lobula and the medulla on both hemispheres. These neurons showed strong wide-field motion responses with complex onset and offset transients, but only weakly responded to narrow, vertical bars, making them substantially different from LO-PNs.

Finally, neurons also connected from the central brain back to the optic lobes (centrifugal neurons; Fig.10c). The input regions of these neurons in the central brain included the arborization areas of the optic lobe projection areas but covered much wider regions. Lobula centrifugal neurons were weakly sensitive to the direction of progressive and regressive optic flow and are suited to send this information to the inner lobula, a region that did not overlap with the inputs of the LO-PNs. Responses of medulla centrifugal neurons were more reminiscent of those of ME-ME-PNs but substantially weaker for all stimuli. They likely feed information back to wide areas of the inner medulla, as well as to layers overlapping with inputs of ME-ME-PNs, potentially allowing for direct feedback loops.

**Table 1:** Data availability. * m = male, f = female, r = right hemisphere, l = left hemisphere; ** no physiology

| Neuron type | IBDB accession number (NIN) | Figure |
| --- | --- | --- |
| LO-PN-bilat-1 | NIN-0000064 (m,l)*, NIN-0000266 (f,l), NIN-0000090 (f,r)* | 2,3,4 |
| LO-PN-bilat-2 | NIN-0000457 (m,l) | 2,3 |
| LO-PN-bilat-3 | NIN-0000256 (f,r) | 2,3,4 |
| LO-PN-bilat-4 | NIN-0000257 (f,l) | 2,3,4 |
| LO-PN-bilat-5 | NIN-0000458** (f,l) | 2 |
| LO-PN-ipsi-1 | NIN-0000224 (f,l;f,r) | 2,3,4 |
| LO-PN-contra-1 | NIN-0000459 (f,l) | 2,3,4 |
| ME-ME-PN-1 | NIN-0000264 (m,l) | 5 |
| ME-ME-PN-2 | NIN-0000269 (f,r), NIN-0000265 (m,l) | 5,6 |
| ME-ME-PN-3 | NIN-0000462 (m,r;f,r) | 5,6 |
| ME-ME-PN-4 | NIN-0000463 (f,r;m,r), NIN-0000254 (f,l) | 5,6 |
| ME-ME-PN-bilat | NIN-0000461 (f,l) | 5,6 |
| MELO-MELO-PN | NIN-0000259 (f,l), NIN-0000255 (f,r) | 7 |
| LO-CN-1 | NIN-0000281 (f,l) | 8 |
| LO-CN-2 | NIN-0000465 (f,l) | 8 |
| ME-CN-1 | NIN-0000262 (f,r) | 9 |
| ME-CN-2 | NIN-0000263 (f,l) | 9 |
| ME-CN-3 | NIN-0000268 (f,l) | 9 |

## Discussion

In the present study we aimed at identifying neurons upstream of the recently identified input neurons to the central complex noduli that are suited to encode the translational velocity of bees during flight (Stone et al., 2017). In particular, we searched for neurons with characteristics that could contribute to the following features of nodulus input neurons: receptive fields covering the entire azimuth, antagonistic motion tuning between right and left visual fields, speed sensitivity, very strong motion responses, lack of transient onset response, and an optimal point of expansion offset by 45° from the animal’s body axis. Our recordings have indeed yielded numerous types of projection neurons from the bee optic lobes that respond to moving wide-field gratings as well as a moving bar. These cell types form three parallel pathways from the lobula and medulla towards large, mostly overlapping regions of the posterior protocerebrum. These regions partly coincided with the dendrites of the noduli input neurons and might thus contribute to generating the complex properties of these cells.

### Complex properties of CX optic flow neurons cannot directly emerge from the identified optic lobe outputs

The responses of one group of lobula projection neurons (group 1 LO-PNs) showed strong, direction-selective responses to grating stimuli, with receptive fields centered in the ipsilateral visual field. The strength of the responses, the clarity of the receptive fields resulting from mapping with a narrow vertical bar, and their pronounced direction selectivity made these neurons the cell types that resembled noduli input cells (TN cells) most closely. As the arborizations of group 1 LO-PNs partially overlapped with the dendritic fields of TN cells, we asked whether the response properties of TN cells could be explained by convergence of several types of the newly found LO-PNs (Fig.10). Despite the superficial similarity in responses, combining the receptive fields of these neurons could not fully recreate the complex tuning properties of TN neurons. TN neurons show no (TN2) or weak responses (TN1) to yaw rotational optic flow (Stone et al., 2017), suggesting that contralateral inhibitory input must balance excitatory ipsilateral input of the same motion direction, or vice versa. This characteristic could be produced, for example, by assuming that LO-PN-ipsi-1 and LO-PN-contra-1 converge on the same downstream neuron with synapses of opposite signs (Fig.10f). However, the receptive fields of all newly found neurons were pointing into anterior directions, extending laterally, but not posteriorly, thus not covering the panoramic receptive fields of TN neurons. Whereas lobula projection neurons with posterior receptive fields might exist, and a combination of inhibitory and excitatory connections could generate a downstream response that is tuned to translational optic flow, further modifications of the response patterns would be needed to shift the preferred center of expansion by 45° away from the midline, as observed in TN neurons. Additionally, the strong transient peaks in response to lights-on switches typical for lobula projection neurons are not found in TN neurons (Stone et al., 2017), indicating that significant low-pass filtering must occur between these two sets of neurons, which is unlikely compatible with a direct connection.

If the presented neurons are involved in relaying optic flow information to TN neurons, we therefore conclude that additional neurons with response characteristics intermediate between LO-PNs (group 1) and noduli input neurons must exist in the bee central brain. Additionally, lobula projection neurons with posteriorly pointing receptive fields are required to provide input to the panoramic receptive fields of TN cells. Given the limited pool of neurons observable through random intracellular recordings, we also cannot exclude that TN neurons receive their input from a different, yet undiscovered pathway.

Indeed, while the similarity of responses is suggestive, our data do not conclusively demonstrate any direct or indirect synaptic connection between lobula outputs and TN neuron input fibers. In the *Drosophila* connectome (hemibrain dataset, Hulse et al., 2021), no LNO or GLNO neuron (the equivalents of bee TN neurons) can be identified that receive synapses from neurons with inputs in the lobula or lobula plate. Although the hemibrain dataset only contains one brain hemisphere, thus missing potential contralateral inputs, it matches our conclusion that there are no direct ipsilateral connections between LO-PNs and noduli input cells in bees. In *Drosophila*, all inputs to LNO cells that are not recurrent from the central complex, result from either local neurons in the lateral accessory lobes, or neurons in the ventrolateral and ventromedial protocerebrum. As the latter regions are served by the bee LO-PNs, the possibility of indirect connections between LO-PNs and noduli input cells in bees is consistent with *Drosophila* connectome data.

### Wide-field motion signals in the optic lobes across insects

As LO-PNs are not receiving direct input from photoreceptor terminals, additional neurons are clearly required to complete the optic flow pathway between the bee retina and the noduli. This is consistent with the established principles of how wide-field motion signals are computed in the insect brain (Shinomiya et al., 2022). In principle, perception of motion is based on the comparison of changes in light intensities at different points in the visual field over time (Cheng and Frye, 2020, Egelhaaf, 2006). Following detection of light by photoreceptors of the compound eyes, extraction of local motion information is performed by retinotopically organized, elementary motion detectors in the medulla and lobula. Local motion is then integrated by large tangential neurons in the lobula plate, thus becoming tuned to wide-field optic flow (reviewed in Borst et al., 2020, Shinomiya et al., 2022). As the lobula complex in bees consists of only one large neuropil, the bee lobula also contains all neurons that are found in the fly lobula plate. The emergence of wide-field motion tuning in the lobula plate is thus consistent with our finding that only neurons of the *Megalopta* lobula responded strongly to wide-field motion and were sensitive to the movement direction and movement speed of gratings. In contrast, neurons of the medulla showed strong responses to a moving vertical bar but were mostly direction-insensitive and gratings only induced weak responses, suggesting that they receive inputs from early stages of motion processing, which is not yet organized in a direction opponent way. Interestingly, direction selectivity was also not present in layer 2 of the lobula, which houses the not direction-selective MELO-MELO-PNs. This suggests that it might be lobula layers 4 and 5, in which direction-selective wide-field motion signals are generated in the *Megalopta* brain.

More broadly, different types of lobula tangential neurons with similar response properties are known from several insect species, e.g. the honey bee (*Apis mellifera*, DeVoe et al., 1982, Hertel and Maronde, 1987, Hertel et al., 1987, Ibbotson, 1991), the hummingbird hawkmoth (*Macroglossum stellatarum*, Wicklein and Varjú, 1999), the bumble bee (*Bombus impatiens*, Paulk et al., 2009a, 2008), the locust (*Locusta migratoria*, Rind, 1990, 2002), a dragonfly (*Hemicordulia spp*., Evans et al., 2019), *and a mantis (Tenodera aridifolia*, Yamawaki, 2018). Lobula plate tangential cells (LPTCs) of flies are most extensively studied in the fruit fly (*Drosophila melanogaster*, Boergens et al., 2018, Schnell et al., 2010, Wei et al., 2020), and the blowfly (*Calliphora vicina* and *C. erythrocephala*, Hausen et al., 1980, Hausen, 1982, 1984). Different types of LPTCs get input from neurons of the medulla and lobula plate (T4 neurons) or the lobula and lobula plate (T5 neurons). These cells are also direction-selective but encode local motion (reviewed in Borst et al., 2020, Shinomiya et al., 2022). As a population, T4 and T5 neurons encode the six optic flow patterns generated by rotational and translational movements (three degrees of freedom each) (Henning et al., 2022). These patterns are transmitted to LPTCs in distinct layers of the lobula plate, providing spatially segregated input. Our finding that LO-PNs (group 1) with opposite direction tuning profiles innervated distinct, adjacent layers of the bee lobula indicates that a similar, layer-specific encoding of directional information could also exist in bees.

### Optic flow signals for non navigational tasks

Our data in *Megalopta* show that not all optic lobe projection neurons have characteristics that make them suited as upstream elements for optic flow based speed sensing in the central complex. This shows that motion cues, including optic flow responses, are required by many downstream circuits. This is in line with optic flow being used as input for many different behaviors. Wide-field motion is naturally generated by movement of objects and scenes in the environment and can be induced by self-motion. It therefore provides information about the 3-dimensional layout of the environment as well as the insect’s self-motion (Egelhaaf, 2006). Thus, besides distance measurement in the context of path integration, optic flow provides key sensory information for different behaviors, including flight control and gaze stabilization using optomotor reflexes (Baird et al., 2011, Huston and Krapp, 2008, Lecoeur et al., 2019, Mauss and Borst, 2020), tracking of prey or the pursuit of mating partners against a natural background (reviewed in Nordström, 2012), as well as collision avoidance (reviewed in Serres and Ruffier, 2017). Of those, best understood are the pathways underlying flight control reflexes, which consist of direct connections between the LPTCs and downstream descending neurons. These cells contact each other in the dorso-lateral deutocerebrum, providing a monosynaptic link to the thoracic ganglia, directly impacting the control of flight-, neck-, and leg-muscles (Mauss and Borst, 2020, Strausfeld and Bassemir, 1985, Strausfeld and Gronenberg, 1990). Similar direct connections were demonstrated for motion-sensitive lobula neurons and descending neurons in the locust (Rind, 1990).

However, the majority of lobula output is not mediated by tangential neurons but by columnar neurons, which receive input from single columns of the lobula that process information from specific, small parts of the environment. In flies, these cells project into the so called optic glomeruli of the protocerebrum (Strausfeld and Okamura, 2007, Wu et al., 2016) that are only sparsely innervated by descending neurons (Okamura and Strausfeld, 2007, Paulk et al., 2009a, Strausfeld and Okamura, 2007). Rather, innervation of these glomeruli by local interneurons and other neurons of the central brain indicate that additional levels of visual processing are localized within the lateral protocerebrum (Okamura and Strausfeld, 2007). Different types of columnar lobula neurons encode distinct visual features and transmit this information into cell-type specific optic glomeruli, forming downstream circuits that have been associated with distinct behavioral modules, e.g. initiation of jumps or backwards walking (Klapoetke et al., 2022, Wu et al., 2016).

While the corresponding neurons also exist in other insects, their output regions are not organized into segregated optic glomeruli but more loosely into contiguous projection fields. In bumblebees they have been shown to mingle with tangential type neurons, in principle similar to our group 2 LO-PNs (Paulk et al., 2009a), and unlike LPTCs, not directly linked to descending pathways. A distinct segregation of projections was present in bumblebees, where tangential neurons of the proximal lobula projected prominently into the superior lateral protocerebrum or the inferior lateral protocerebrum, whereas neurons from distal lobula layers and large-field medulla neurons predominantly arborized in the posterior protocerebrum. This anatomical segregation was accompanied with a functional segregation into predominantly motion-sensitive neurons in the distal lobula layers and color-sensitive neurons in the proximal lobula (Paulk et al., 2009a, 2008). This is consistent with our data from *Megalopta*, in which all neurons that encoded motion stimuli were located in the posterior protocerebrum. However, as there is no comprehensive, functionally grounded map of optic lobe projections in any insect to which we could compare our *Megalopta* neurons, and the number of visually guided behaviors is large in highly visual insects like bees, it is not possible at this point to associate our recorded neurons with any specific functional pathway or behavior, besides the already described potential links to speed sensing in the context of path integration.

### Conserved patterns and species-specific aspects

We finally asked whether any of the neurons and pathways in the current study could possibly be specific to *Megalopta*. When comparing the recorded *Megalopta* neurons to optic lobe projection neurons of other insects, it is difficult to find specific neuron counterparts for several reasons. First, the variable morphology of the lobula complex with different numbers of subdivisions (from one in hymenoptera, two in most orders, to up to six in dragonflies or mantids, Fabian et al., 2020, Rosner et al., 2017) leads to divergent morphologies of the input regions of LO-PNs, predicting different neural shapes even for homologous neurons. Second, the different layout of the target regions in the central brain leads to a similar problem. While the same super-regions (according to Ito et al., 2014) exist across the brains of all insects, their detailed shape, borders and size differ markedly between orders and species. Finally, the physiological properties are difficult to compare, as recordings have been accumulated since the 1970s, with widely varying stimulus paradigms. Therefore, comparisons can only be done at a high level.

If we compare our recorded neurons to other species in broad categories, we find mostly conserved patterns of information flow with no obvious specializations in *Megalopta*. Tangential LO-PNs in flies, honeybees, and other species generally project to posterior brain regions with different cell types serving ipsilateral, contralateral and bilateral regions (Hausen, 1984, Hertel et al., 1987). direction-selective motion responses are typically found in these cells. In contrast, across species, tangential neurons associated with the medulla generally did not show direction selectivity (Rind, 1987) and were often characterized by spectral responses (Hertel et al., 1987, Paulk et al., 2009b).

The segregation into weakly and strongly motion-sensitive neurons we found in *Megalopta* has not been reported in other insects. However, due to the lack of comparable data, it is currently unclear whether the projection fields in the *Megalopta* central brain associated with these different types of motion responses are a consistent feature of insect brains. Similarly, little is known about neurons that provide feedback to the optic lobe. Whereas medulla neurons with centrally placed somata were reported in the honeybee, their strong responses to ipsilateral light flashes suggests that they are actually medulla output cells (Hertel et al., 1987). However, LPTCs in *Drosophila* are known to change their motion tuning during walking behavior (Chiappe et al., 2010) and flight (Maimon et al., 2010) and the activity of LPTCs in *Calliphora vicina* is influenced by haltere oscillations that indicate enhanced motor activity (Rosner et al., 2010). These observations demonstrate that non-visual information about the locomotor state of the fly enters the optic lobe. The lobula centrifugal neurons reported in the current study might be suited to relay such information to the bee lobula, in particular as they only marginally responded to visual stimuli. Finally, the recorded medulla centrifugal neurons physiologically resembled a weak version of the inter-medulla projection neurons and also terminated in matching layers of the medulla. This provides a potential substrate for a direct recurrent connection between the medulla and the central brain, that might fine-tune the responses of inter-medulla projection neurons in response to non visual stimuli. Similar neurons that provide feedback to the medulla from the ocelli and the posterior protocerebrum have been recently reported in bees and were implicated in color constancy (Garcia et al., 2017).

Lastly, having performed all recordings in a nocturnal species inhabiting dense jungle environments, raised the question whether there are links between neural responses and the nocturnal lifestyle of *Megalopta*. While the principle response of neurons was qualitatively independent of light levels across a considerable range, we were not able to test the sensitivity threshold of motion responses. This was mostly due to the difficulty in identifying motion-sensitive neurons with very dim stimuli, and the long time that would be needed to dark adapt animals in which neurons were identified with bright stimuli. As recordings did not last longer then 3-5 minutes on average, dark adapting during ongoing recordings was not feasible. Nevertheless, we observed that the majority of neurons responding to wide-field motion saturated at stimulus velocities between 90-100°/s. If these neurons are truly tuned to speed rather than temporal frequencies, as suggested by our data, it is interesting that the saturating stimulus velocity corresponds to the preferred speed of optic flow experienced by *Megalopta genalis* in a flight tunnel (Baird et al., 2011). This suggests that these neurons might be optimized for processing optic flow during the slow flight speed typical for this species. Similar behavioral data in bumblebees shows that these insects fly faster (Baird et al., 2011), predicting that corresponding neurons should be tuned to higher stimulus velocities. Indeed, recordings from wide-field motion stimuli across hoverflies, hawkmoths and bumblebees revealed that the velocity optimally driving these neurons was matched to lifestyle: around 200°/s in fast flying bumblebees, and 50°/s in both hovering species (O’Carroll et al., 1996). Similar data from dragonflies yielded a value of 135°/s (Evans et al., 2019).

Overall, our data show that general patterns of information flow in the context of motion vision are conserved between *Megalopta*, bumblebees, and flies, groups of insects that are separated by hundreds of millions of years of evolution. Yet, as most projection areas of optic lobe output neurons are located in poorly understood brain regions, it is difficult to link specific neurons to concrete behavioral tasks that are not reflex-driven. The likely speed encoding by translational optic flow during path integration offers an exceptionally clear link to navigation behavior and our results provide a starting point towards understanding how general purpose optic flow output from the lobula of the optic lobe is being shaped to meet the specific requirements of path integration.

## Acknowledgments

The authors wish to thank Dr. Julia Schuckel for help in collecting *Megalopta* and Ola Gustafsson for assistance in confocal microscopy. The authors are grateful for financial support from the following organizations: the Swedish Research Council (Vetenskapsrådet; 621-2012-2213 to S.H., and FA8655-07-C-4011 to E.W.), the European Research Council (ERC) under the European Unions Horizon 2020 research and innovation program (grant agreement no. 714599 to S.H.), a Marie-Curie Intra-European Fellowship (327901 to S.H.), the Air Force Office for Scientific Research (2012-02205 to E.W.) and the Wenner-Gren Foundation (to A.H.) and a donation from Jennifer and Greg Johnson (to W.W.).

## Author Contributions

A.H. and S.H. performed all experiments; A.H., R.H., A.A. and S.H. analyzed the data; R.H. wrote the first draft of the manuscript together with S.H.; R.H., K.K., and S.H. designed the figures; E.W. and W.W. provided guidance, equipment, field access, and helped conceiving the study together with S.H.; all authors co-wrote and reviewed the final version of the manuscript.

## Supplementary Data

**Figure S1:**
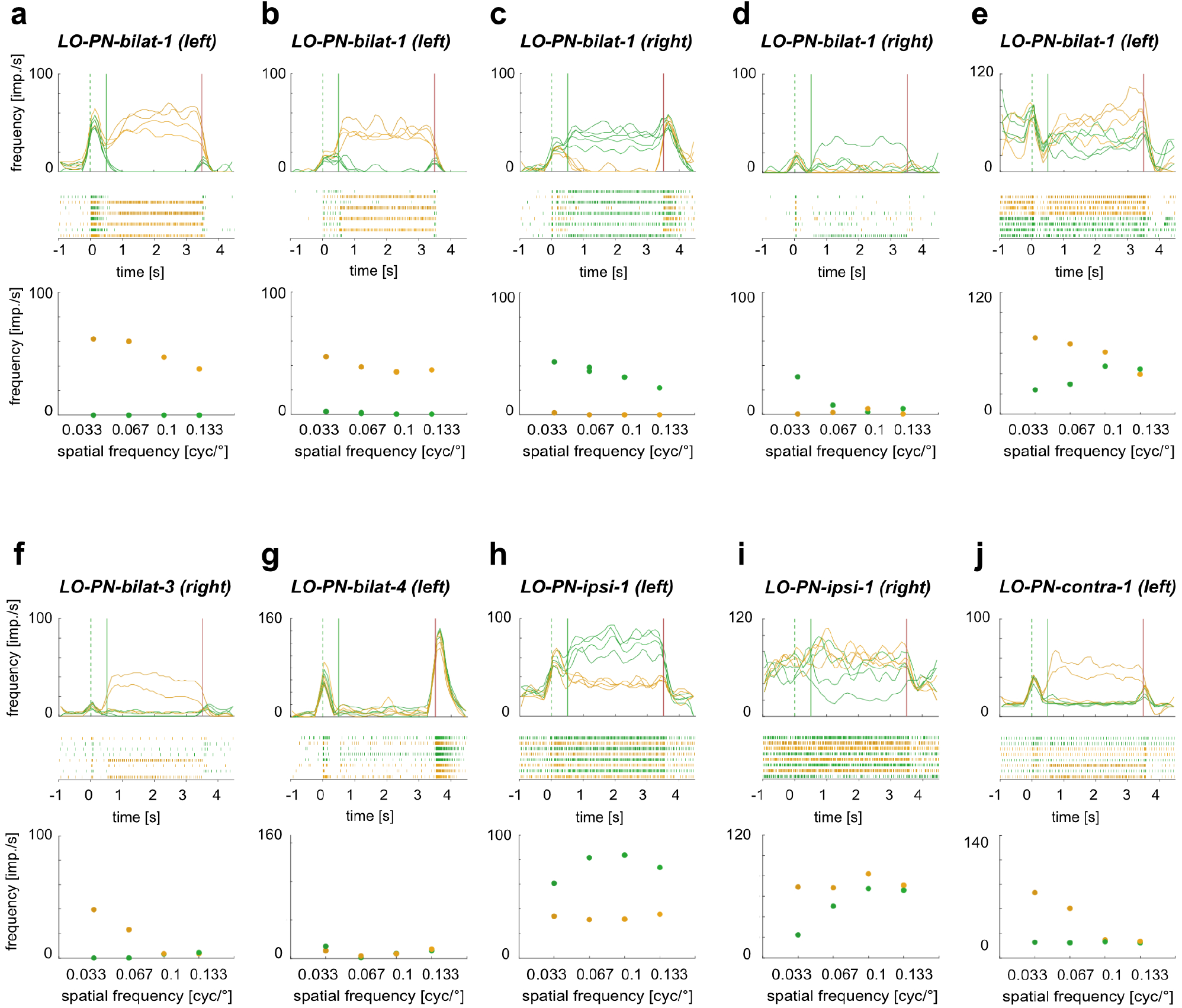
Spatial frequency tuning of lobula projection neurons (LO-PNs). (a-j) Top: Activity of different types of LO-PNs in response to clockwise (yellow) and counter-clockwise (green) wide-field optic flow of different spatial frequencies (for spatial frequency values see bottom graph). Green dotted lines: grating presented; green solid lines: motion onset; red lines: motion stop. Middle: Corresponding raster plots. Bottom: Mean frequency during the final 2 s of each stimulus bout. No ND filter in (a-c,e,h).

**Figure S2:**
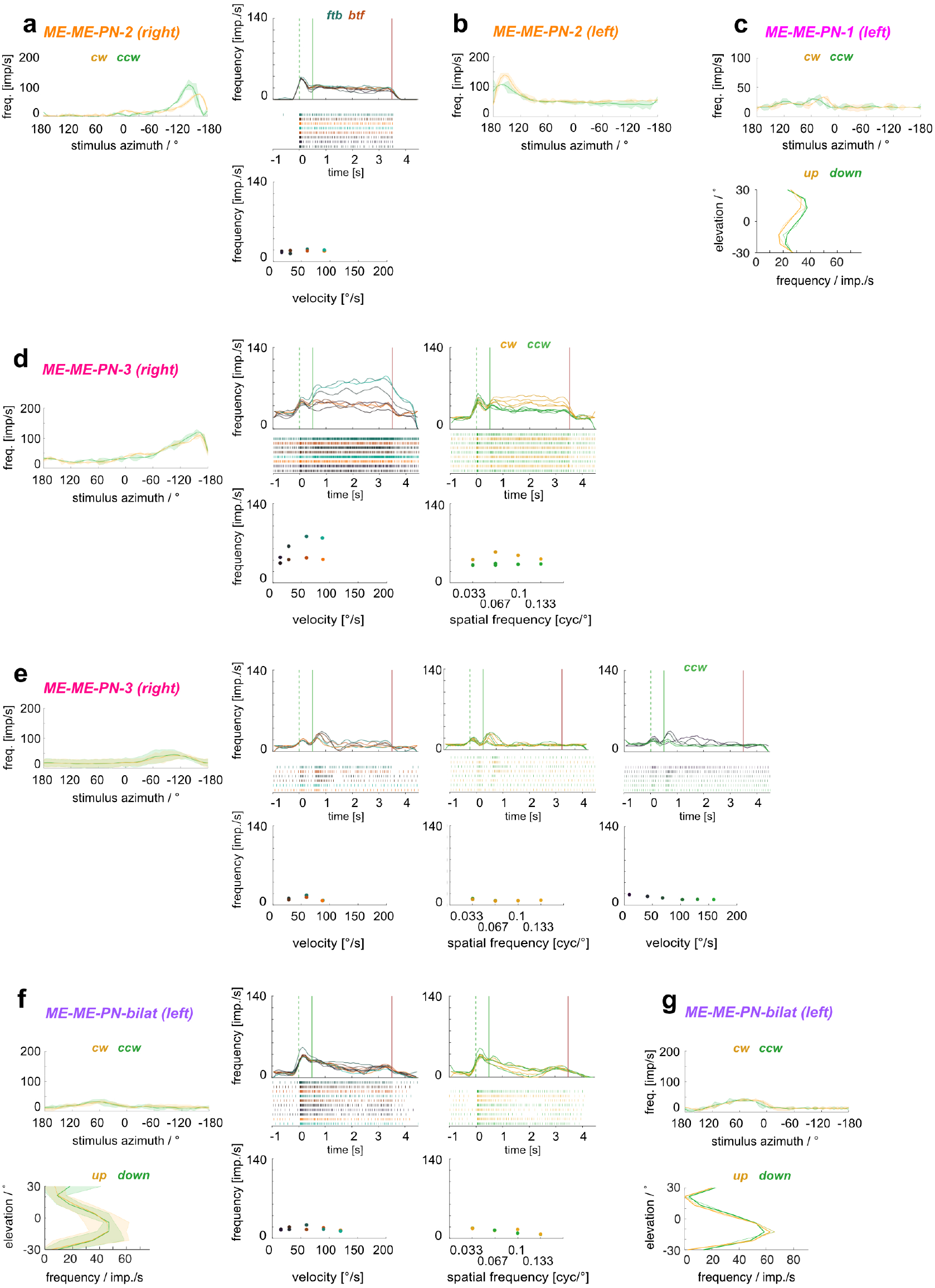
Physiological data for all ME-ME-PN-1,-2,-3, and -bilat neurons. Horizontal receptive fields of all cells (a,d,e: left; b; c,g: top; f: top left corner) were mapped with a green vertical bar moving around the animal either clockwise (cw, yellow) or counter-clockwise (ccw, green). Vertical receptive fields of three cells (c,g, bottom; f, bottom left corner) were mapped with a horizontal bar moving up (yellow) or down (green). Activity in response to front-to-back (ftb) or back-to-front (btf) optic flow of different velocities was recorded in one ME-ME-PN-2 (a, right graphs), both ME-ME-PN-3 (d,e) and one ME-ME-PN-bilat (f). Activity in response to cw and ccw optic flow of different spatial frequencies was recorded in both ME-ME-PN-3 cells (d,e) and in one ME-ME-PN-bilat cell (f). Activity in response to ccw optic flow of different velocities was recorded in one ME-ME-PN-3 cell (e, graphs on the right). No ND filter in (d,e), one ND filter in (g).

**Figure S3:**
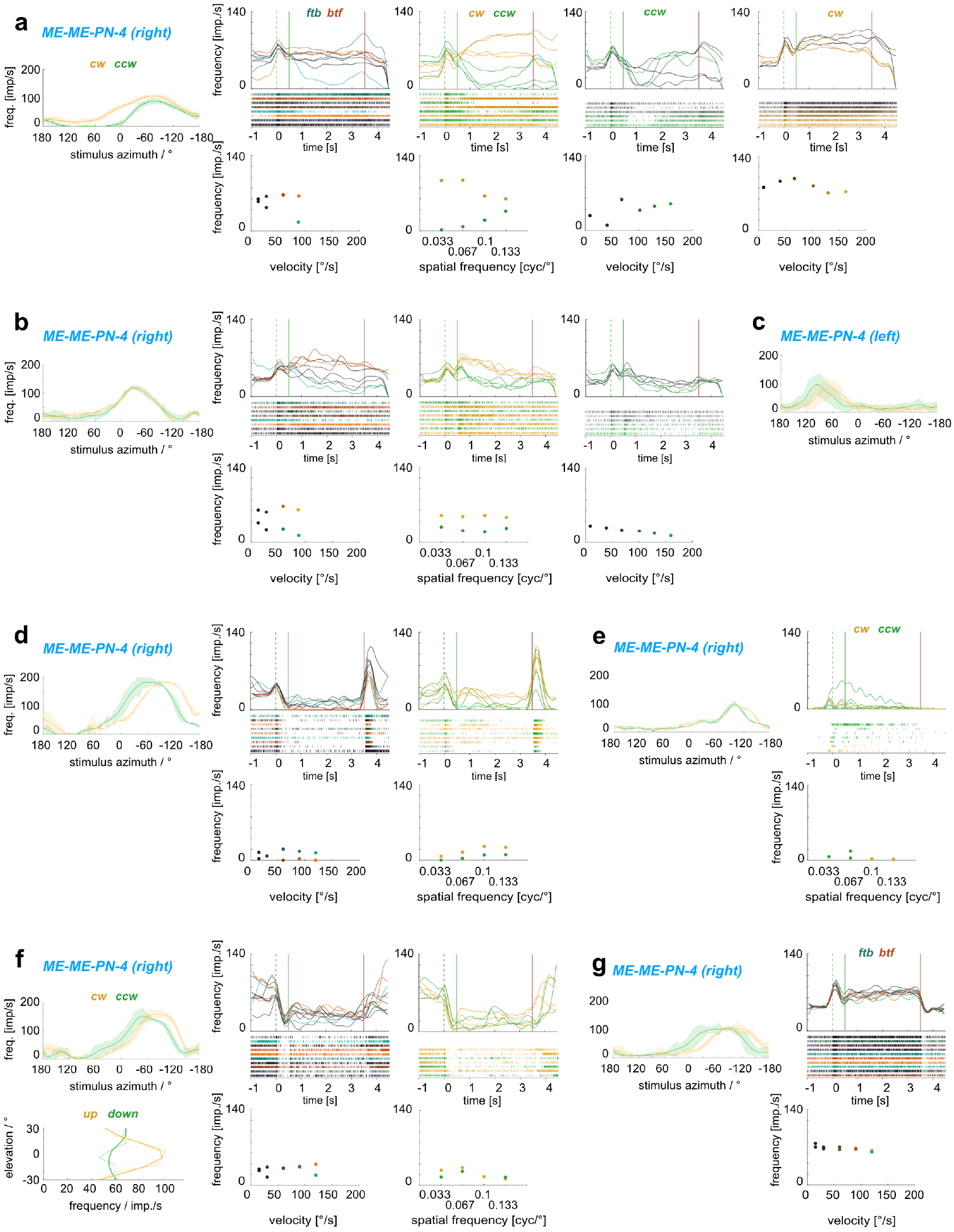
Physiological data for all ME-ME-PN-4 neurons. Physiological data for all ME-ME-PN-4 neurons. Horizontal receptive fields of all cells (a,b,d-g: left; c) were mapped with a green vertical bar moving around the animal either clockwise (cw, yellow) or counter-clockwise (ccw, green). Vertical receptive fields of one cell (f, bottom left corner) was mapped with a horizontal bar moving up (yellow) or down (green). Activity in response to front-to-back (ftb) or back-to-front (btf) optic flow of different velocities was recorded in five out of seven cells (a,b,d,f,g). Activity in response to cw and ccw optic flow of different spatial frequencies was recorded four cells (a, b, d, f). Activity in response to cw and ccw optic flow of different velocities was recorded in one cell (a). Activity in response to ccw optic flow of different velocities was recorder in one cell (b). No ND filter in (a,b,d,e).

